# Confidence-modulated shared control for reliable intracortical BCI navigation in dynamic environments

**DOI:** 10.64898/2026.03.23.713738

**Authors:** Ophelie Saussus, Pinhao Song, Sofie De Schrijver, Irene Caprara, Thomas Decramer, Renaud Detry, Peter Janssen

**Affiliations:** Laboratory for Neuro- and Psychophysiology, Department of Neurosciences, KU Leuven and the Leuven Brain Institute, Leuven, Belgium; Department of Mechanical Engineering, Research unit Robotics, Automation and Mechatronics, KU Leuven, Leuven, Belgium; Department of Electrical C Computer Engineering, University of Washington, Seattle, WA, USA; Research group Experimental Neurosurgery and Neuroanatomy, KU Leuven and the Leuven Brain Institute, Leuven, Belgium; Department of Electrical Engineering, Research unit Processing Speech and Images, KU Leuven, Leuven, Belgium

## Abstract

Brain–computer interfaces enable people with paralysis to control neuroprosthetic devices, but reliable operation in dynamic environments remains limited by noise in decoded neural commands. We developed a confidence-modulated probabilistic shared-control framework that adaptively integrates the decoded neural commands with a probabilistic temporal prior to stabilize execution while preserving the directional structure of the decoded command. In two male macaques performing closed-loop tasks in complex environments, shared control nearly eliminated execution-level failures, while maintaining the directional structure of neural commands. Abrupt target changes revealed a limitation: temporal-history dependence transiently slowed responsiveness when recent movement history became inconsistent with the updated command, although this delay was not obligatory. Resetting the temporal prior removed the residual shared-control delay and restored performance, demonstrating a specific contribution of temporal history to the impairment. Here, we show that shared control improves the reliability of continuous intracortical brain-computer interfaces while revealing a trade-off between stabilization and responsiveness.

## Introduction

Brain–computer interfaces (BCIs) aim to restore movement by translating neural activity into control signals for external devices. Both invasive and non-invasive systems now achieve impressive performance in controlled laboratory settings, enabling cursor control, constrained reaching, and basic manipulation ^1,2^. Continuous intracortical BCI control has also been demonstrated in two-dimensional ‘maze’ tasks requiring curved trajectories around obstacles ^3^, as well as in our previous virtual-navigation paradigm ^4^. In these studies, obstacle avoidance was achieved under direct neural control rather than through a downstream shared controller. However, clinical adoption will require these systems to operate reliably beyond constrained laboratory settings, where assistive devices must move through cluttered and changing environments while preserving the user’s evolving intended actions.

A major challenge for real-world BCI control is that decoded neural commands can be noisy. During continuous control, short-timescale fluctuations in decoded velocity may not reflect genuine changes in user intent, but they are nevertheless expressed directly in device movements. When such commands are used to control physical assistive devices such as powered wheelchairs or robotic manipulators, this variability could create safety risks, including collisions or unintended device actions ^5^. Reliable BCI use therefore requires more than accurate decoding of movement intent: it requires mechanisms that stabilize the execution of decoded commands while remaining responsive to meaningful changes in intent.

One strategy is to combine neural control with shared-control assistance that stabilizes unreliable commands and compensates for environmental constraints ^6,7^. Shared-control systems can use information about the environment to improve safety and reduce the burden of continuous control. However, assigning too much control authority to the assistive system risks overriding the user’s intended actions and may reduce the sense of agency that is central to the acceptance and long-term use of assistive neurotechnology ^8,9^. This creates a central design problem for shared-control BCIs ^7,10^: assistance must correct unreliable execution without substituting the user’s decisions.

Shared control has been explored in both non-invasive and invasive BCIs. In many non-invasive BCI systems, decoded commands are low-dimensional or discrete, and reliable navigation often requires substantial autonomous support ^6–8,10–13^. By contrast, invasive BCIs can provide continuously varying control signals that contain rich information about intended movement. In this regime, the goal of assistance is not to determine the trajectory autonomously, but to stabilize noisy commands when intent is present yet unreliable. While shared-control elements have been incorporated into invasive systems for robotic reaching, grasping, and manipulation to improve robustness^5,14–20^, previous approaches have often relied on fixed blending or phase-specific assistance, with limited focus on continuously adapting assistance to fluctuations in decoded control reliability. Consequently, confidence-modulated continuous shared control for high-bandwidth invasive BCI navigation under competing demands for stabilization and rapid responsiveness has not previously been demonstrated.

Here, we develop a confidence-modulated probabilistic shared-control framework for invasive BCI navigation under dynamic environmental constraints and abrupt changes in intended goals. The controller integrates decoded neural velocity commands with a probabilistic model of recent movement history and environmental constraints, allowing it to stabilize execution while preserving the directional structure of the decoded command. We test this framework in two macaques performing closed-loop virtual navigation using intracortical BCI control (Saussus et al., 2026^4^). The tasks impose complementary demands on the controller: obstacle-avoidance tasks required stabilization of noisy but goal-consistent commands, whereas a target-respawn task required rapid updating after an abrupt change in goal. We show that shared control markedly improves navigation reliability by suppressing execution-level failures such as collisions and target overshoot, while leaving the directional structure of the decoded command largely intact. Rapid goal changes reveal a limitation of temporal stabilization, in which assistance can transiently delay responsiveness when recent action history is no longer predictive. Together, these results identify confidence-modulated shared control as a general post-decoder strategy for improving the safety and reliability of continuous invasive BCI control, while defining design constraints for future neuroprosthetic systems operating in dynamic environments.

## Results

To evaluate how shared control arbitrates between stabilization and responsiveness under decoder ambiguity, we examined three navigation tasks imposing opposing demands on the controller in two nonhuman primates implanted with three 96-channel Utah arrays (Blackrock Neurotech) (**Figure 1a**). Neural activity was recorded from primary motor cortex (M1), dorsal premotor cortex (PMd), and ventral premotor cortex (PMv), with 189 ± 6 electrodes contributing to the decoder in Monkey 1 and 151 ± 11 in Monkey 2. During online control, decoded neural velocities either directly controlled the virtual sphere (BCI-only trials) or were passed through the shared-control controller before execution (shared-control trials; **Figure 1b**).

**Figure 1.**
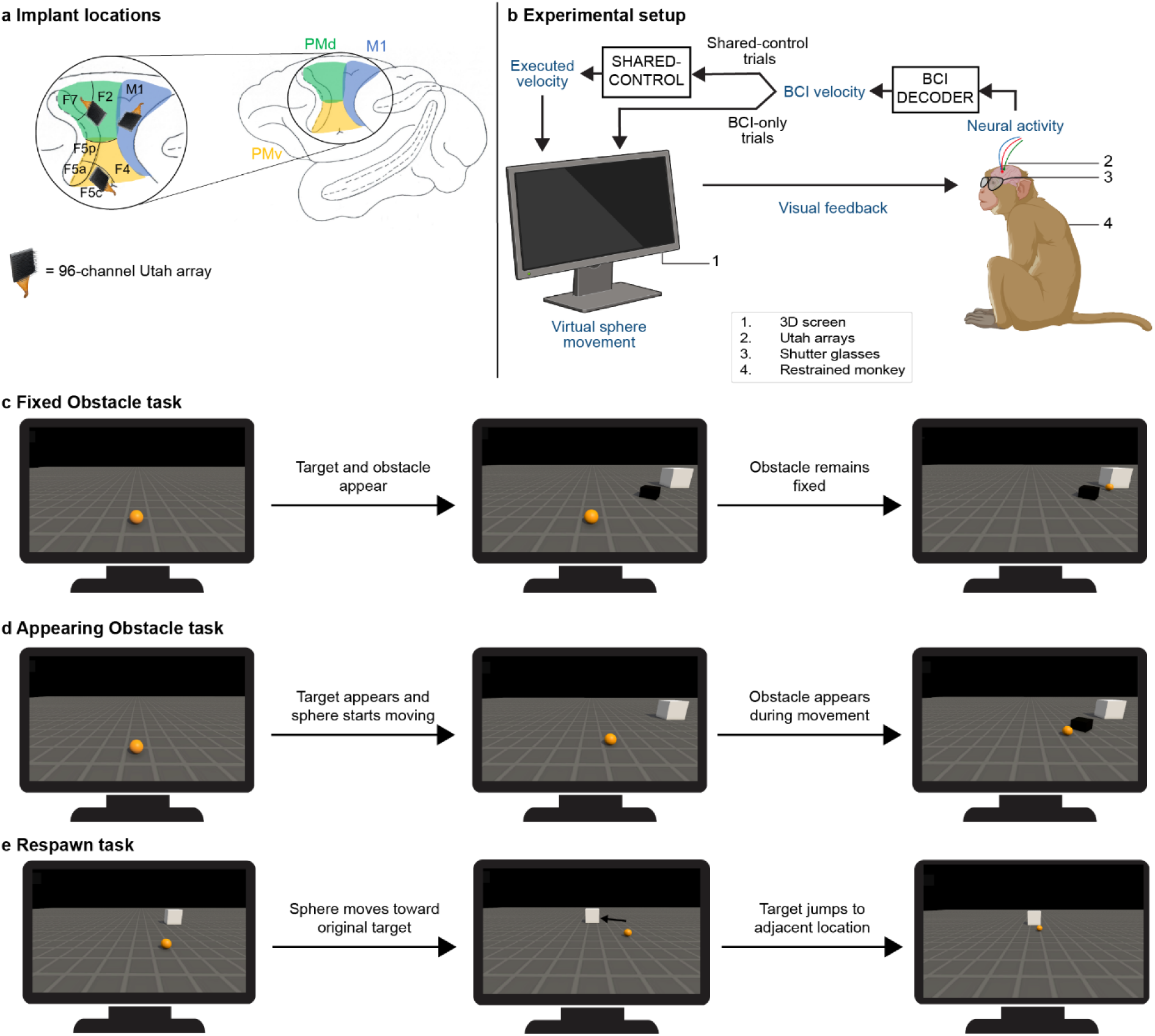
Experimental setup and task paradigms. **a**, Schematic of the cortical implant locations common to both monkeys. Each monkey was implanted with three SC-channel Utah arrays targeting primary motor cortex (M1), dorsal premotor cortex (PMd), and ventral premotor cortex (PMv). **b**, Closed-loop experimental setup. Neural activity was decoded in real time to generate brain-computer interface (BCI) velocity commands. In BCI-only trials, decoded velocities directly controlled movement of the virtual sphere. In shared-control trials, decoded velocities were passed through the shared-control controller, which generated the executed velocity. Movement of the virtual sphere was displayed on the 3D screen, providing continuous visual feedback to the monkey. **c**, Fixed Obstacle task. The target and obstacle were present from trial onset, and the obstacle remained fixed throughout the trial. **d**, Appearing Obstacle task. The sphere initially moved toward a fixed target before an obstacle appeared unexpectedly during movement. **e**, Respawn task. The sphere initially moved toward the cued target, which subsequently changed to an adjacent target location during movement. Figure adapted from Saussus et al., Science Advances, 202C (CC BY 4.0).

In the first task, decoded velocities from the intracortical BCI controlled a sphere navigating toward a fixed target while avoiding a static obstacle positioned between the start location and the goal (Fixed Obstacle; **Figure 1c**). This task, previously performed under neural control alone^4^, provided a benchmark for assessing whether shared-control stabilizes execution under stable environmental structure. In a second task, the obstacle emerged unexpectedly during the trial (Appearing Obstacle; **Figure 1d**), allowing us to test stabilization under sudden environmental perturbations while the intended goal remained unchanged. In a third task, the target location was reassigned mid-trial (Respawn; **Figure 1e**), requiring rapid abandonment of prior action history and responsiveness to an abrupt change in goal intention.

These tasks impose complementary demands: obstacle tasks require suppressing execution variability, whereas the Respawn task requires rapid updating to goal changes. Demonstrating effective arbitration across both regimes provides a stringent test of adaptive shared-control under ambiguity in the decoded control signal. Representative trials appear in Supplementary Movies 1–3.

### Probabilistic shared-control framework

To mitigate noise in the decoded velocity, we implemented the shared-control framework Robot Trajectron V2 (RT-V2)^21^ as a post-processor to the neural decoder (**Figure 2**a). The controller operated at 20 Hz (50 ms updates) during closed-loop control. The shared-control prior *p*(*ξ* ∣ ℎ*_t_*, *c*) represented candidate future velocity trajectories ξ, conditioned on recent movement history ℎ*_t_* (last eight 50 ms steps, 400 ms total) and environmental context *c*. At each prediction step, the prior action distribution was represented as a velocity-space Gaussian mixture with mode-specific weights, mean velocities, and covariances, from which candidate future trajectories were generated (see Methods for the mathematical formulation). The environmental context comprised the candidate target and obstacle locations; the controller received the locations of all five candidate targets but was not provided with the identity of the target cued to the monkey. The instantaneous decoded BCI velocity *v* was not incorporated when computing the prior.

**Figure 2.**
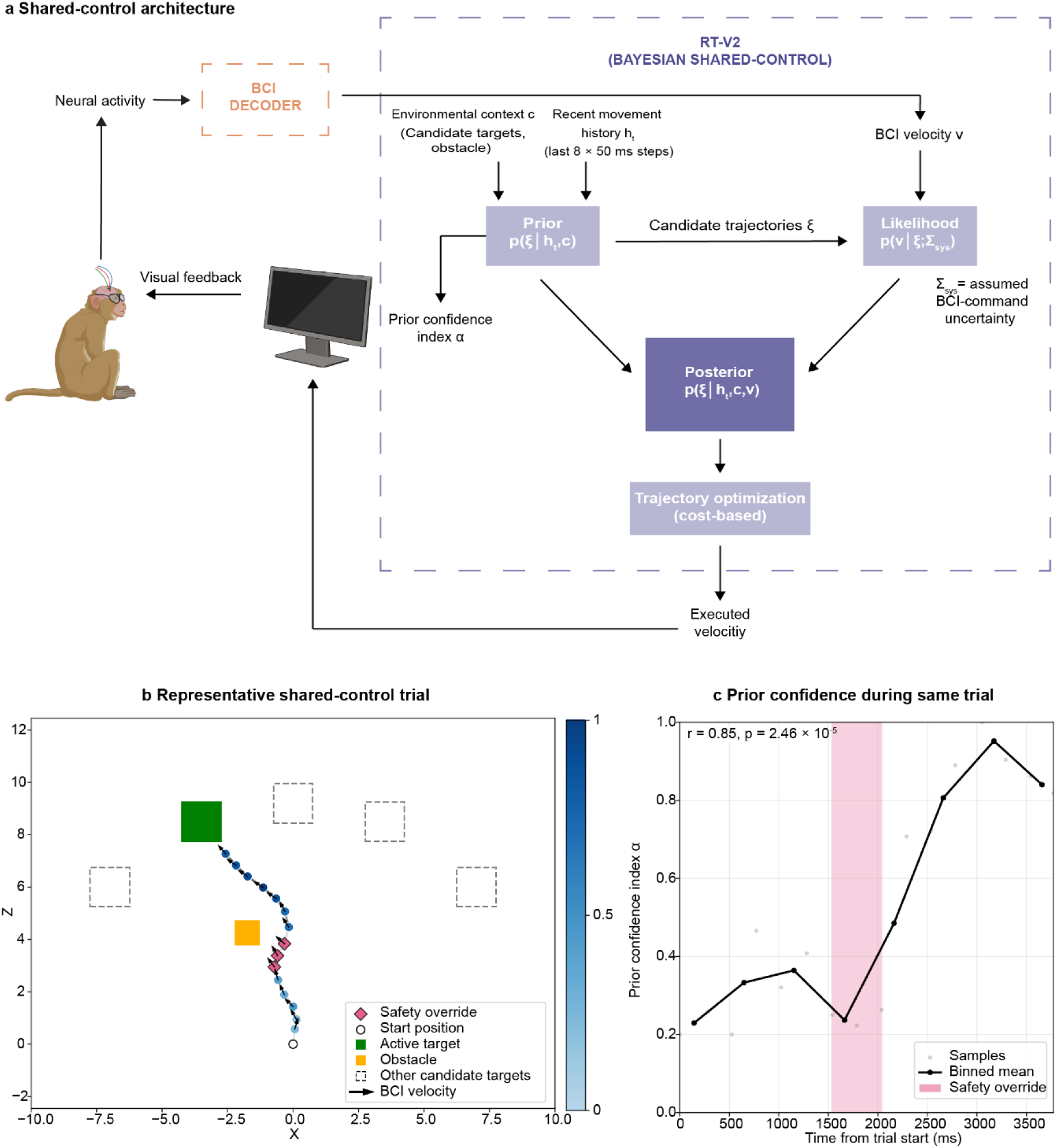
Probabilistic shared-control architecture and representative closed-loop trial. **a**, Shared-control architecture. Neural activity was decoded every 50 ms to produce a two-dimensional brain-computer interface (BCI) velocity v. Robot Trajectron V2 (RT-V2) generated a multimodal trajectory prior p(ξ ∣ ℎ_t_, c) from recent movement history ℎ_t_ and environmental context c, comprising candidate target and obstacle locations. The prior action distribution was parameterized as a velocity-space Gaussian mixture, *p(v_t_ | h_t,c_) = Σ_z_π_z_N(v_t_; μ^z^_t_,Σ^z^_t_*, from which candidate future trajectories were generated. The instantaneous decoded BCI velocity was incorporated through the Gaussian likelihood p(v ∣ ξ; Z_sys_), with Z_sys_ specifying the assumed uncertainty in the decoded BCI command. Combining the prior and likelihood yielded a posterior distribution over candidate trajectories, which were evaluated according to task and safety costs; the instantaneous velocity of the selected trajectory was then executed. The prior confidence index α was defined as *clip_[0.2,1]_(1 − H^LB^_t_/H_max_)*, where *H^LB^_t_* is the entropy lower bound of the prior action distribution and H_max_ = log(K_est_), with K_est_ estimated independently for each trial from the maximum observed entropy. α_t_ ranged from 0.2 to 1, with larger values indicating a more concentrated prior. Identical α values can produce different executed-velocity corrections depending on the instantaneous BCI command, its relationship to the prior, and Z_sys_. RT-V2 received all candidate target and obstacle locations but was not provided with the identity of the target cued to the monkey. The trajectory model was pretrained on simulated navigation trajectories and transferred to online BCI control without fine-tuning on monkey BCI trajectories. **b**, Representative shared-control trial from the Fixed Obstacle task (Monkey 2). The open circle indicates the start position, the green square the cued target, the orange square the obstacle, and dashed squares the remaining candidate targets. Black arrows show decoded BCI velocities; blue points show the executed trajectory, coloured by α; magenta diamonds indicate safety-override samples. **c**, Prior confidence index α during the same trial. Grey points show individual samples, the black line shows the binned mean, and shading indicates the safety override. During an imminent-collision override, the instantaneous BCI command was excluded for three consecutive 50-ms updates (150 ms) before standard shared control resumed.

During online control, the decoded velocity *v* was incorporated through a Gaussian likelihood *p*(*v* ∣ *ξ*; Σ_sys_), which assigned higher probability to candidate trajectories whose instantaneous velocity was more consistent with the decoded BCI command. The decoded velocity was modeled as a noisy observation of the intended velocity, with Σ_sys_ specifying the assumed uncertainty in the BCI command. The posterior trajectory distribution was then obtained using Bayes’ rule (Equation 1).

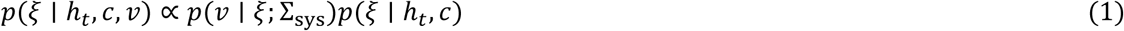

Candidate trajectories were sampled from this posterior and evaluated using a cost function that penalized obstacle collisions and rewarded progress toward candidate targets. The selected trajectory determined the instantaneous executed velocity.

We quantified the concentration of the trajectory prior using the prior confidence index *α*, derived from the normalized entropy lower bound of the prior action distribution (**Figure 2**a; see Methods for the exact definition). High *α* indicates a more concentrated prior, whereas low *α* indicates greater uncertainty. Importantly, *α* is a derived measure of prior confidence rather than an explicit blending coefficient. Its value therefore does not uniquely determine the correction applied to the decoded command; this depends on the interaction between the prior, the instantaneous BCI likelihood and Σ_sys_. The trajectory model was pretrained on simulated navigation trajectories and transferred to the online monkey BCI task without fine-tuning on monkey BCI data. For invasive BCI implementation, the continuously decoded two-dimensional BCI velocity served as RT-V2’s user-command input. Σ_sys_ was selected empirically during pilot testing and subsequently held fixed, and an imminent-collision safeguard temporarily excluded the BCI command for three consecutive updates (150 ms). The underlying probabilistic shared-control architecture was otherwise unchanged.

To illustrate the resulting closed-loop behavior, **Figure 2**b,c shows a representative navigation trial from the Fixed Obstacle task with shared control enabled (Monkey 2). In Error! Reference source not found.**b**, black arrows indicate the decoded BCI velocity, while blue dots show the executed sphere trajectory colored by the prior confidence index, where higher α indicates a more peaked prior and therefore greater confidence in the prior’s predicted action distribution. Early in the trial, α is low (**Error! Reference source not found.c**) and the executed path closely follows the decoded BCI command, consistent with relatively weak constraint from the prior. As the prior becomes more concentrated and α increases, the executed trajectory becomes progressively smoother and more directly target-oriented while remaining consistent with the decoded BCI command.

When the current decoded BCI velocity would lead to an imminent collision, the controller triggers a brief safety override lasting three consecutive 50 ms updates (150 ms), during which the instantaneous BCI command is temporarily excluded and the executed velocity is selected from the shared-control prior (magenta diamonds in Error! Reference source not found.**B**; shaded interval in **Error! Reference source not found.c**). In this example, the decoded BCI velocity pointed directly toward the obstacle, illustrating that continuous velocity commands can be locally unsafe in cluttered environments. The override redirects the sphere around the obstacle, after which the controller returns to the standard shared-control regime. Across all shared-control trials, explicit safety override intervals occupied 9.7% of total movement time. Per trial, the median cumulative override exposure was 3.7 equivalents of a 150 ms override interval (IQR 0.3–5.7), corresponding to a median cumulative duration of approximately 0.54 s. Time-pooled exposure was 6.7% in Fixed Obstacle and 12.1% in Appearing Obstacle; the Appearing Obstacle aggregate includes the six-month follow-up sessions.

The representative trial suggested that decoded input continued to shape control as the movement progressed. We tested this directly in closed-loop simulations in the same BCI-like five-target obstacle environment. Incorporating the instantaneous user command changed controller output throughout the movement: mean posterior–prior action differences were 0.98, 0.42, and 0.41 m/s in the early, middle, and late phases, respectively, with corresponding cosine alignment to the user command of 0.77, 0.84, and 0.91 (all *p* < 10^−66^). Replacing the target-directed command after 67% projected progress strongly redirected subsequent movement (alignment: 0.11 baseline vs. 0.72 perturbed; paired difference, +0.61; 95% CI, +0.49 to +0.73; *p* = 1.7 *x* 10^−17^) and produced a mean post-activation trajectory separation of 1.23 m (95% CI, 1.13–1.33 m; *p* = 5.76 *x* 10^−44^). Thus, ongoing user input continued to influence the controller even late in movement.

### Shared-control improves reliability of BCI navigation

Task success, defined as reaching the cued target and remaining within the target window for 500 ms, was significantly higher with shared-control across all obstacle-navigation tasks, with consistent gains across subjects and stable performance over months (**Figure 3**). In the Fixed Obstacle task (**Figure 3a**), shared-control substantially improved performance in both monkeys, yielding ∼30 percentage-point (pp) increase in success rates compared to BCI-only control. This improvement was significant in both animals (paired Wilcoxon signed-rank (two-sided): Monkey 1: median Δ = 31.3pp, *p* = 4.88 × 10^-4^; Monkey 2: median Δ = 26.8pp, p = 9.77 x 10^-4^). Monkey 2, who was naïve to this task at the start of the experiments, showed a gradual improvement across sessions even in the absence of assistance (Spearman’s ρ = 0.755, p = 0.01), whereas Monkey 1—previously trained on this task^4^—reached an average success rate of approximately 80% with shared-control, matching its earlier No-Obstacle benchmark performance (BCI-only).

**Figure 3.**
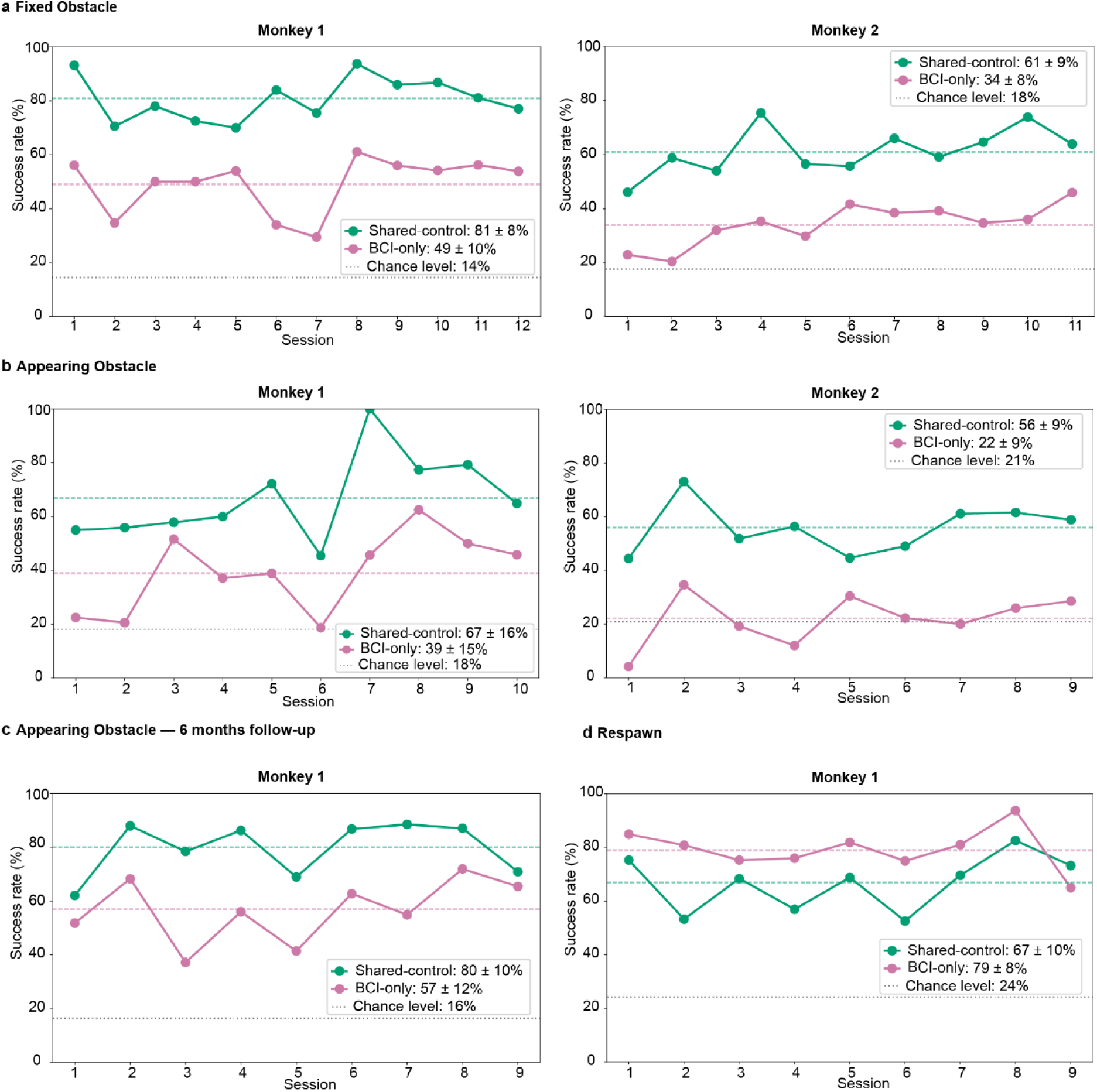
Success rates across tasks and sessions for shared-control and BCI-only. Success rates per session for Monkey 1 (left) and Monkey 2 (right). Green lines show shared-control trials, pink lines show brain-computer interface (BCI)-only trials. Dashed lines indicate average success rates, and gray dashed lines indicate chance levels. Text annotations within each panel report the across-session mean ± standard deviation success rate for shared-control and BCI-only conditions. Shared-control improves performance in the obstacle-based tasks (A–C), whereas performance is reduced in the Respawn task (D). **a,** Fixed Obstacle task. **b,** Appearing Obstacle task (December). **c,** Appearing Obstacle task (C months later, Monkey 1). **d,** Respawn task. The number of sessions (N) for each task is provided in Extended data Table 2.

Performance gains with shared-control persisted in the more challenging Appearing Obstacle task (**Figure 3b**), in which an obstacle appeared unexpectedly during navigation. In this task, shared-control produced large improvements in success rates (Monkey 1: 67% vs. 39%; median Δ = 26.7pp, p = 1.95 × 10^-3^; Monkey 2: 56% vs. 22%; median Δ = 35.6pp, p = 3.91 × 10^-3^). Without assistance, Monkey 2’s performance remained near chance, underscoring that shared control was essential for successful navigation under these conditions. The increase in success was accompanied by a modest speed–safety trade-off. On successful trials, path-length efficiency did not differ, whereas shared control increased total acquisition time (Appearing Obstacle: +0.68 s; Fixed Obstacle: +0.58 s) and reduced mean movement speed (Appearing Obstacle: 1.84 vs. 2.19 m/s, *p* = 3.81 × 10^−6^; Fixed Obstacle: 1.97 vs. 2.30 m/s, *p* = 2.38 × 10^−7^). At the same time, shared control increased minimum obstacle clearance by approximately 0.48 Unity units and reduced mean squared jerk (Appearing Obstacle: 3.55 × 10^5^vs. 5.04 × 10^5^, *p* = 2.67 × 10^−4^; Fixed Obstacle: 3.76 × 10^5^ vs. 5.27 × 10^5^, *p* = 1.67 × 10^−5^), indicating slower but smoother and safer successful movements (**Extended Data Figure 1; Extended data Table 1**).

To assess the stability of shared-control benefits over time, we repeated the Appearing Obstacle task in Monkey 1 six months later, with no additional training or practice on this task during the intervening period (**Figure 3c**). Average BCI-only performance was higher in the follow-up than across the original experiment (57% vs. 39%); however, performance at the beginning of the follow-up was similar to that reached at the end of the original experiment (final three original sessions: 52.8%; first three follow-up sessions: 52.4%; **Figure 3**b,c). Importantly, the shared-control benefit remained substantial after removal of the hidden fixed obstacle that had been present in the original sessions (mean gain: 27.4 vs. 23.0 percentage points*; difference = 4.4 percentage points, 95% confidence interval (CI) −5.8 to 15.0; p = 0.5SC*). Shared control increased performance to approximately 80%—comparable to both the Fixed Obstacle task and prior No-Obstacle benchmarks under BCI-only control.

In contrast to obstacle-based tasks, the Respawn task—in which the target location changed mid-trial (**Figure 3d**)—imposed an opposing demand on the controller by requiring rapid updating of previously consistent action history. Under these conditions, shared-control reduced performance relative to BCI-only control (Monkey 1: 67% vs. 79%; median Δ = −11.4pp, p = 0.01), consistent with a mismatch between the shared-control module’s temporal assumptions and abrupt changes in the decoded control signal. This result highlights a limitation of shared-control when abrupt goal changes invalidate the recent action history underlying the controller’s temporal prior.

### Comparison with alternative shared-control strategies

To determine whether the performance of our confidence-modulated shared-control framework could be reproduced by simpler forms of assistance, we performed matched closed-loop simulations comparing full shared control with fixed blending, prior-only control, and hard-coded phase-specific assistance (Extended Data Tables 3–4; Methods). We tested these strategies in two environments: a BCI-like five-target obstacle environment with one obstacle and a randomized cluttered environment with 1–6 candidate targets and 1–6 static obstacles. Fixed blending combined the current user command and trajectory-prior action using a constant weight throughout the trial. In the BCI-like environment, full shared control achieved 97% success compared with 88% for the best tested fixed blend (W=0.75; paired difference, +9 percentage points; 95% CI, +2 to +17; exact McNemar p = 0.035). In the more variable cluttered environment, success was similar (75% vs. 73%), but full shared control produced fewer collisions (4% vs. 19%, *p* = 2.7 *x* 10^−4^) and the best-performing fixed weight differed between environments (W = 0.75 vs. 0.85). Hard-coded phase-specific assistance, which replaced the user command with the trajectory-prior action whenever the sphere came within a predefined distance of an obstacle, performed near ceiling for several distance thresholds in the structured environment but was more sensitive to this predefined distance in the cluttered environment, where success ranged from 42% to 73%.

Prior-only control retained environmental context and recent movement history but excluded the instantaneous user command. It achieved only 28% success in the BCI-like environment and 17% in the cluttered environment, indicating that the trajectory prior alone was insufficient to reproduce the performance obtained when the current user command was incorporated. To determine whether this poor performance reflected an inability to navigate or an inability to identify the intended target, we performed a separate target-only simulation without obstacles. With a single candidate target, prior-only control reached it on 82% of trials, with the remaining 18% timing out. With five candidate targets, prior-only control reached any target on 83% of trials but the designated target on only 14%, compared with 100% and 84%, respectively, for full shared control. Across 1–20 candidate targets, correct-target success declined from 100% to 44% for full shared control and from 82% to 2% for prior-only control, while prior-only any-target success remained 91% with 20 candidates. Thus, the prior could frequently generate plausible goal-directed movements but became increasingly unable to identify the intended target as the number of candidate goals increased.

Because the shared-control controller includes an explicit safety override for imminent collision, we next asked how much of the observed shared-control benefit could be explained by collision prevention alone. Perfect rescue of every BCI-only “blocked at obstacle” failure accounted for 17.4 of the 28.7 percentage-point gain in the Appearing Obstacle task and 13.2 of the 29.4 percentage-point gain in the Fixed Obstacle task, leaving residual gains of 11.3 and 16.2 percentage points, respectively. Because the benefit of shared control may depend on the reliability of the incoming command, we also tested robustness to increasing directional variability in the simulated user command in the randomized cluttered environment. Across angular-error standard deviations of 0°, 10°, 20°, 40°, and 60°, user-only success was 43%, 42%, 44%, 38%, and 33%, respectively, whereas full shared control maintained success rates of 88%, 90%, 90%, 87%, and 80%. Thus, the shared-control benefit remained between 45 and 49 percentage points across the tested range of user-command reliability, while collision incidence under full shared control remained at 1%. Together, these control analyses showed that simpler strategies could perform well under specific conditions, but none reproduced the overall performance profile of full shared control across environments.

### Shared-control benefit depends on baseline BCI performance

We next examined how the benefit of shared-control depended on baseline BCI performance across target directions, pooling data across sessions, monkeys, and tasks (**Figure 4**). Although shared-control generally improved success rates, the magnitude of improvement varied with baseline (BCI-only) performance. Across the 255 target × session observations, shared-control gain showed significant negative curvature with baseline BCI performance (session-clustered *p* = 1.45 × 10^−6^; **Figure 4**c). Notably, target × session observations with exactly 0% BCI-only success were based on relatively few BCI-only trials (median 5 trials) but still showed positive directional alignment of the decoded command with the cued target (mean trial-level cosine similarity = 0.49 across 208 trials; >0–10% baseline observations: 0.47 across 116 trials), indicating that zero behavioral success did not imply absence of directional information in the decoded command. Binned averages showed the largest gains at low-to-intermediate baseline performance, with progressively smaller gains as baseline performance approached ceiling. Because the two monkeys sampled different ranges of baseline performance, we repeated this analysis separately by monkey and task (**Extended Data Figure 2**). Significant negative curvature was observed overall in both monkeys and within the Fixed Obstacle task in both monkeys. The separate fits for the original Appearing Obstacle datasets and the Monkey 1 six-month follow-up did not reach statistical significance. A model allowing the relationship to differ between monkeys provided a better fit than a common-curve model (likelihood-ratio test, *p* = 0.018), indicating some between-monkey variation in the shape of the relationship. Cross-validated prediction errors were nevertheless similar between the common and monkey-specific models. Thus, the pooled relationship was not solely driven by between-monkey differences, but its precise shape varied across subjects and tasks.

**Figure 4.**
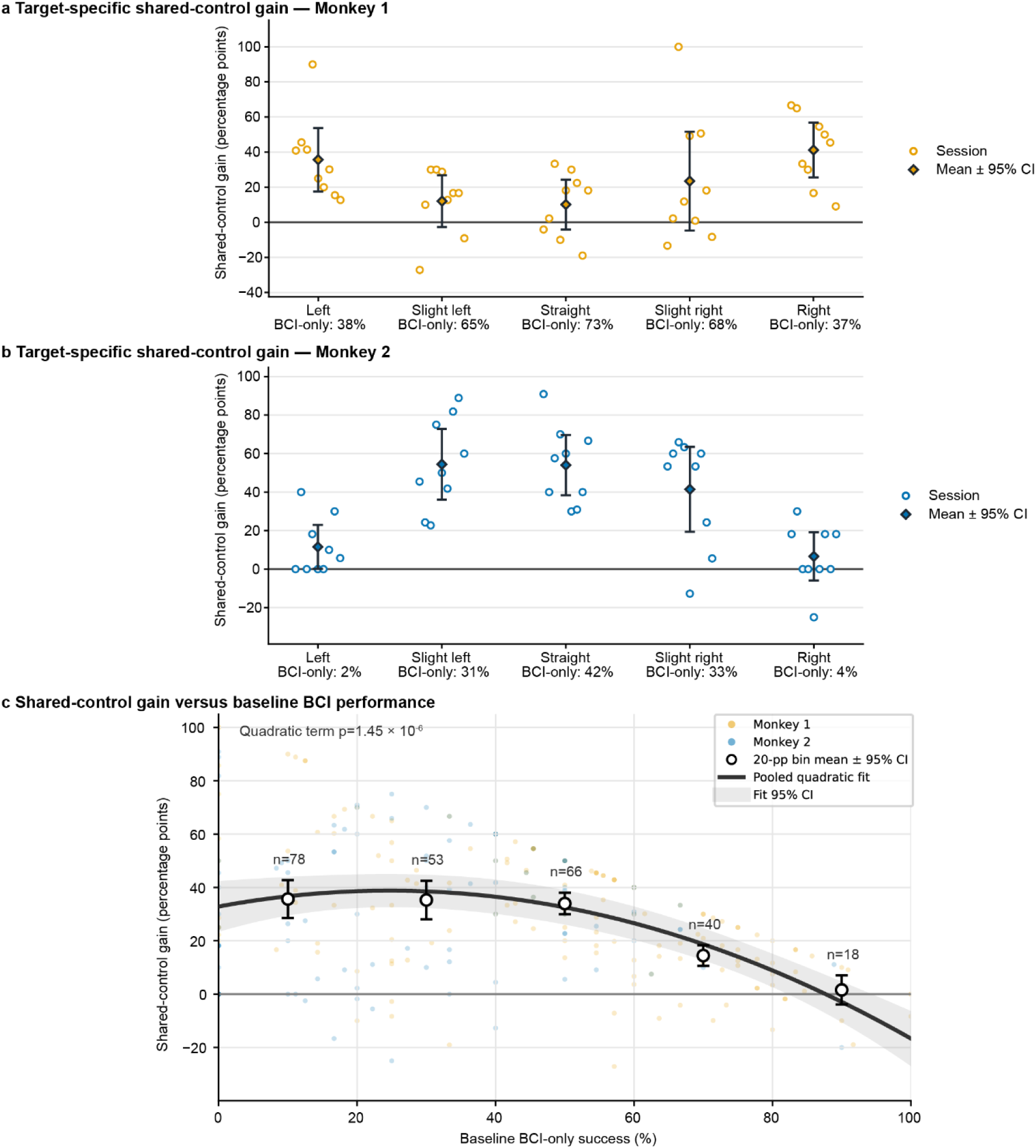
Per-target effects of shared-control. **a**, Monkey 1 – Appearing Obstacle (6months later): mean per-target shared-control gain in percentage points (pp) (diamonds and error bars indicate the mean and 95% confidence interval across sessions; circles indicate individual sessions) across sessions (n = S), baseline BCI-only success rates shown below each target. **b,** Monkey 2 – Appearing Obstacle: mean per-target shared-control gain across sessions (n = S) (diamonds and error bars indicate the mean and 95% confidence interval across sessions; circles indicate individual sessions), with baseline (BCI-only) success rates. **c,** Relationship between baseline BCI-only success and shared-control gain across all 255 target × session observations from 51 sessions. Faint points show individual observations; open circles and error bars show mean gain and 95% confidence intervals within 20-percentage-point baseline-performance bins, with the number of observations indicated for each bin. The black curve shows the pooled quadratic fit estimated from individual observations, and the shaded band indicates its 95% confidence interval. The model included monkey and task intercepts, with standard errors clustered by session (two-sided z test for the quadratic term, p = 1.45 × 10^−6^).

This relationship is illustrated by per-target examples from each monkey (in the Appearing Obstacle task, **Figure 4a–b**). The two animals sampled complementary portions of the pooled relationship. In Monkey 2, the two targets with the lowest baseline success showed little gain, whereas targets with intermediate baseline performance showed large improvements with shared control. Monkey 1 sampled a generally higher baseline-performance range, with smaller gains for the highest-performing targets. Panel C therefore shows the full analysis comprising 255 target × session observations from 51 sessions, together with binned summaries indicating the number of observations contributing to each baseline-performance range. Together, these analyses indicate that shared-control gain decreased as baseline BCI performance approached ceiling. However, low baseline success did not necessarily indicate an absence of usable decoded information, as even target × session observations with 0% BCI-only success retained positive directional alignment with the cued target. Thus, the ability of shared control to improve poor behavioral performance may depend on the goal-relevant directional information retained in the decoded command.

### Shared-control stabilizes decoded commands

To isolate the effect of shared-control on execution, we examined execution-related failure modes across monkeys and tasks, including trials in which the sphere became blocked at an obstacle, overshot the cued target, failed to approach the target sufficiently closely, or failed to dwell within the target window for the required duration (representative examples in Extended Data Figure 3), expressed as a fraction of all trials (**Figure 5a**). Without shared-control, execution failures were common: ∼16% of trials terminated with the sphere against the obstacle, and a similar fraction overshot the cued target window without satisfying the dwell criterion. With shared-control enabled, execution failures were strongly suppressed, decreasing from ∼37% of BCI-only trials to ∼4%. This reduction reflected elimination of obstacle-related failures and a decrease in target overshoot errors (∼16% to ∼3%; paired Wilcoxon signed-rank test, one-sided, p < 0.05, n=5). Remaining execution-related failure categories were also significantly reduced, each falling to near-zero levels (one-sided paired Wilcoxon, all p < 0.05). Consistent with the controller-comparison analysis above, eliminating “blocked at obstacle” failure alone could not account for the full shared-control benefit, indicating that shared control also reduced other execution-related failures. Together, these results show that shared control robustly stabilizes execution, preventing trajectory-level failures caused by noisy velocity decoding in cluttered environments.

**Figure 5.**
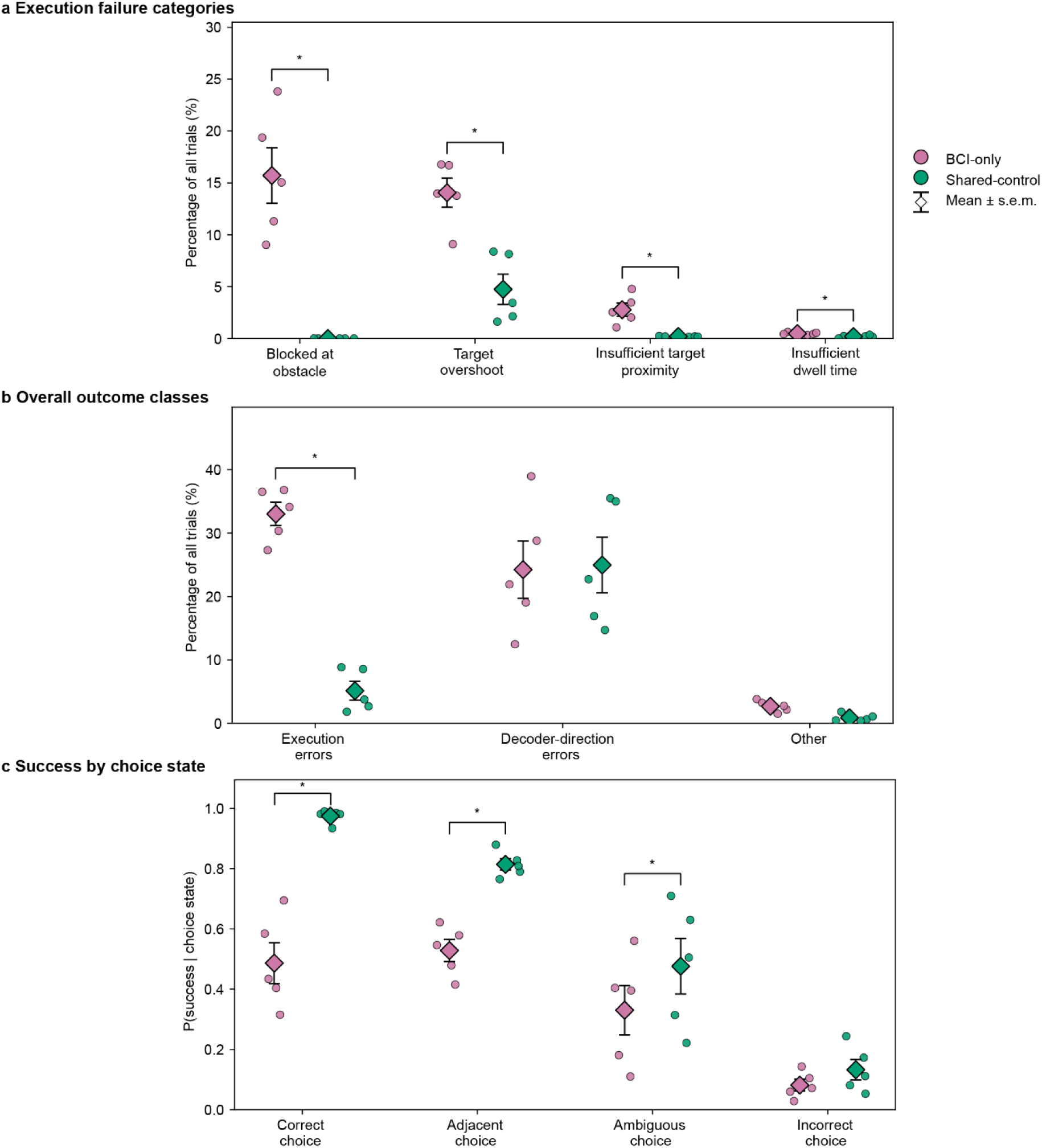
Failure modes and shared-control benefits across monkeys and tasks. **a,** Proportion of trials falling into execution-related failure categories for brain-computer interface (BCI)-only (pink) and shared-control (green), shown across monkey x task combinations. Execution failures include trials in which the sphere became blocked at the obstacle, overshot the cued target window without satisfying the dwell criterion, failed to approach the target closely enough (did not reach the target window), or failed to dwell within the target window for the required 500 ms. **b,** Distribution of trial outcomes grouped into execution errors, decoder-direction errors, and other failures, expressed as a fraction of all trials. Execution errors correspond to the trajectory-level failures shown in panel A. Decoder-direction errors refer to trials in which the decoded BCI velocity was not consistently aligned with the cued target direction and include adjacent, ambiguous, and incorrect decoder-inferred choice states defined in panel C. “Other” pools remaining non-success outcomes. **c,** Success probability conditioned on the decoder-inferred choice state for the same monkey × task units. “Correct” and “Incorrect choice” denote trials in which the cued or a specific non-cued target, respectively, accounted for at least 60% of decoded movement directions. Among the remaining trials, “Adjacent choice” denotes those in which at least 80% of the non-true-target directional mass was assigned to adjacent targets; all others were classified as “Ambiguous choice”.. Asterisks indicate p < 0.05 in paired Wilcoxon signed-rank tests across the five monkey × task units. Tests were one-sided for reductions in panel a failure proportions and panel b execution errors, and for increases in panel c success probabilities. Tests for panel b decoder-direction errors and other failures were two-sided. The number of sessions (N) for each task is provided in Extended data Table 2.

Once execution errors were largely eliminated, the distribution of the remaining trial outcomes shifted (**Figure 5b**). Decoder-direction errors, defined as trials in which the decoded BCI velocity was not consistently aligned with the cued target direction, accounted for a larger fraction of unsuccessful trials under shared-control because execution-related failures were strongly suppressed, rather than because their absolute frequency increased. Other failure modes remained rare in both conditions. As a result, errors in the decoded velocity direction became the dominant remaining limitation on performance. These decoder-based classifications are distinct from non-cued target acquisition, which refers to the behavioral outcome in which the executed trajectory ultimately acquired a non-cued candidate target location.

To relate this redistribution of outcomes to the quality of the decoded command, we next conditioned success probability on the decoder-inferred choice state (**Figure 5c**). When the decoded BCI velocity consistently pointed toward the cued target (“correct choice”), shared-control nearly doubled the probability of success, approaching ceiling performance. Shared-control also significantly improved success when decoded directions were biased toward adjacent targets (“adjacent choice”), although performance remained lower than in the correct-choice regime. Shared-control also significantly improved success when decoded command was ambiguous, though performance remained intermediate. In contrast, when trials were classified as an “incorrect choice” state, meaning that the decoded command predominantly favored a specific non-cued candidate target direction, success probabilities remained low in both conditions.

Together, these results indicate that shared control selectively stabilizes execution without correcting misdirected decoded commands. When the decoder provides at least a partially correct estimate of the cued-target direction, the shared-control controller can suppress execution noise and convert otherwise unstable trials into successful outcomes. By contrast, when the decoded command consistently favored a non-cued target direction, shared-control provided little benefit, leaving misdirected decoded commands as the main limitation of performance. Thus, low baseline task performance alone does not preclude substantial shared-control benefit: assistance can rescue unreliable execution when the decoded command retains at least partial information about the cued-target direction, but provides little benefit when the command consistently favors a specific non-cued target.

### Rapid goal changes reveal history-dependent control limits

We next examined how the prior-confidence index α evolved around abrupt task events and whether its dynamics help explain both performance gains and failures. Because *α* was normalized independently within each trial, it was interpreted as a relative within-trial measure of prior concentration rather than an absolute cross-trial measure of prior confidence, with higher values indicating a more concentrated prior and lower values indicating a broader, more uncertain prior. Moreover, α should not be interpreted as a direct measure of assistance strength: similar α values can correspond to different correction magnitudes depending on the instantaneous decoded BCI command and its relationship to the prior. Across tasks involving unexpected environmental changes, α exhibited a stereotyped, event-locked modulation (Figure **6**). In the Appearing Obstacle task (**Figure 6a**), obstacle onset was followed by a rapid dip in α and a gradual recovery over the subsequent ∼1 s, a pattern that was consistent across monkeys and sessions. This modulation was tightly time-locked to the true obstacle appearance and substantially larger than fluctuations observed around pseudo-events placed at random times along the same trajectories (controlling for slow within-trial trends in α), indicating that it reflects a transient disruption of the controller’s short-term action history rather than slow trial-level trends. As the trajectory stabilized, α recovered to (and often exceeded) its pre-event level, reflecting increasing concentration of the controller’s prior action distribution as recent movement became more consistent.

**Figure 6.**
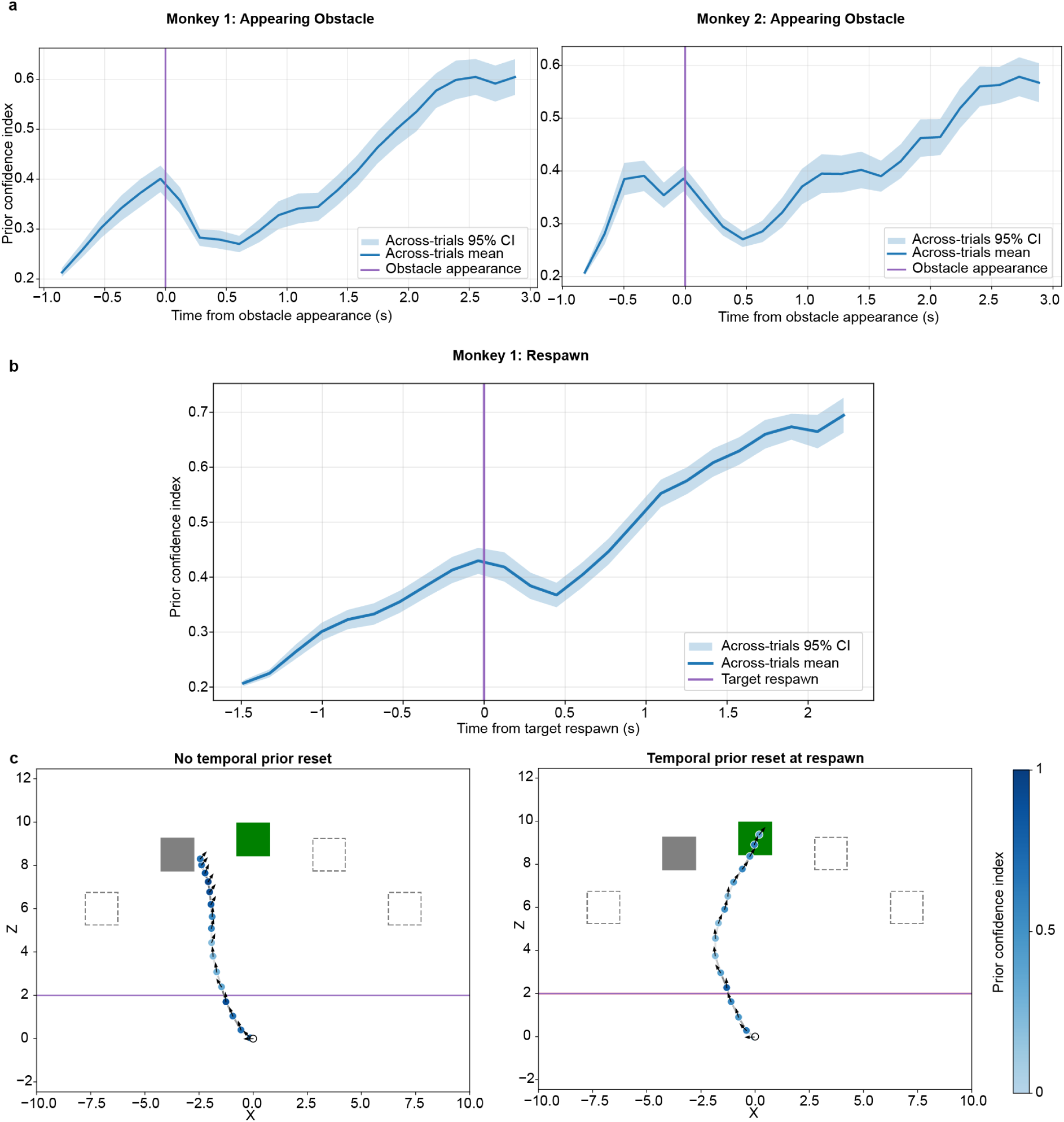
Event-locked modulation of the shared-control temporal prior and its behavioral consequences. **a,** Appearing Obstacle task. Across-trial mean shared-control prior confidence index α (defined as 1 − normalized entropy of the prior action distribution) aligned to obstacle appearance (t = 0 ms, vertical purple line) for Monkey 1 (left) and Monkey 2 (right). **b,** Respawn task for Monkey 1, showing α aligned to target respawn (t = 0 ms). In **a** and **b**, the blue trace denotes the across-trial mean, shaded bands indicate the S5% confidence interval across trials. Confidence bands represent across-trial variability for visualization; statistical comparisons were performed on session-level summaries. Obstacle appearance and target respawn both induce a transient reduction in prior confidence, reffecting an abrupt environmental change and conffict with recent action history, respectively. For the Appearing Obstacle task, the modulation was significantly larger than ffuctuations observed around pseudo-events placed at random times along the same trajectories (Monkey 1: median real–pseudo Δα difference = −0.042, p = 0.011; Monkey 2: −0.040, p = 0.012; two-sided Wilcoxon signed-rank tests). For the Respawn task, the event-aligned modulation also differed significantly from pseudo-event ffuctuations (median real–pseudo Δα difference = −0.01C, p = 0.028). **c,** Same-trial offline replay isolating the effect of the temporal prior. Identical neural data and decoded velocities were replayed with the temporal prior either carried over across respawn (left) or reset at respawn (right). Resetting the prior eliminates the residual shared-control lag and restores baseline performance, demonstrating that temporal inertia in the controller contributes to the impairment. The number of sessions (N) for each task is provided in Extended data Table 2.

A similar dip in α was observed following target respawn events (**Figure 6b**), indicating that the prior-confidence index is sensitive to abrupt violations of the recent movement history on which the controller’s temporal prior is based. In contrast, no comparable modulation was observed in the Fixed Obstacle task (**Extended Data Figure 4**), where environmental constraints were present from trial onset and recent action history remained predictive throughout the trial.

The Respawn task provided a direct test of whether this temporal history necessarily imposed a delay when the intended movement changed abruptly. In the Respawn task, shared-control reduced success relative to BCI-only control (Monkey 1: 67% vs. 79% success). However, such a delay was not obligatory. On successful shared-control trials in which both decoded BCI and executed velocities reoriented toward the new target, the median BCI-to-executed reorientation lag was 0 ms, indicating that the executed command could follow the updated BCI command within the same 50 ms sampling interval despite the pre-jump movement history.

Failures were instead associated partly with the incoming decoded BCI command itself. Decoded BCI velocity reoriented toward the new target less reliably on unsuccessful than successful trials both with shared control (75.4% vs. 99.8% within 2 s) and during BCI-only control (73.3% vs. 100.0%). Among trials in which the decoded BCI command did reorient, reorientation was also delayed on unsuccessful trials in both conditions (shared control: 683.0 vs. 384.3 ms, Δ = +298.7 ms, *p* = 1.95 × 10^−21^; BCI-only: 543.0 vs. 449.8 ms, Δ = +93.2 ms, *p* = 0.00975). In shared-control trials, these differences persisted across analysis windows from 1–3 s. The BCI-only results also indicated that delayed or absent updating of the decoded BCI command was associated with Respawn failure even in the absence of the shared-control mechanism.

Shared control nevertheless contributed an additional temporal delay on unsuccessful trials. Among trials in which both the decoded BCI and executed velocities reoriented, the median BCI-to-executed reorientation lag was 59.3 ms on unsuccessful trials compared with 0 ms on successful trials (*p* = 7.0 × 10^−5^), and the controller required longer to abandon the pre-jump action history (median Δt_decay = +289 ms, q = 0.020). Offline replay of the same trials with the temporal prior reset at the moment of respawn eliminated this residual lag and restored performance to approximately the BCI-only level (∼80%), demonstrating a specific contribution of the controller’s short-term temporal prior to the shared-control impairment (**Figure 6c**).

Together, these results show that temporal integration does not impose an inevitable responsiveness cost. The same shared-control architecture could follow an updated BCI command without measurable additional delay on successful trials, whereas failures reflected both slower or absent updating of the decoded BCI command and an additional delay associated with the controller’s recent action history. Thus, temporal history created the potential for inertia after abrupt goal changes, but the occurrence of delayed control was also associated with the quality and timing of the incoming decoded BCI command rather than following inevitably from the shared-control architecture.

## Discussion

Reliable and safe control remains a central barrier for invasive brain–computer interfaces, particularly in dynamic environments where variability in decoded neural signals can produce unstable or unintended device behavior. Here we show that confidence-modulated shared control improves the reliability of continuous intracortical BCI navigation by stabilizing how decoded commands are executed. This stabilization produced large and consistent gains in obstacle-based tasks by suppressing execution-level failures, whereas abrupt target changes revealed a potential cost of temporal integration: recent action history could contribute to delayed responsiveness on unsuccessful trials. These asymmetric effects reveal a core design constraint for shared-control BCIs: assistance must stabilize unreliable commands without compromising responsiveness to genuine changes in the decoded command.

A central insight is that shared control improved performance primarily by stabilizing execution, rather than altering the directional content of the decoded command. Across tasks, shared control nearly eliminated collisions and target overshoot, whereas trials in which the decoded velocity consistently favored a non-cued target remained difficult. Conditioning performance on the decoder-inferred choice state confirmed this distinction: assistance was most effective when decoded commands were at least partially aligned with the cued target, but provided little benefit when the decoded command consistently favored a non-cued candidate target direction. Thus, the controller converted noisy but goal-consistent commands into successful trajectories, while leaving misdirected decoded commands largely unchanged. This indicates that shared control acted as an execution-stabilizing layer, not as an autonomous intent-correction system.

The magnitude of shared-control benefit also depended on baseline BCI performance, since shared-control gain decreased as baseline performance approached ceiling. However, low behavioral success did not necessarily indicate an absence of usable decoded information: even cases with 0% BCI-only success retained positive directional alignment with the cued target. Together with the decoder-inferred choice-state analysis, these findings indicate that poor behavioral performance can still be improved when the decoded command retains goal-relevant directional information, whereas assistance provides little benefit when the command consistently favors a specific non-cued target. Behavioral success therefore provides only an indirect measure of decoder quality and does not by itself determine whether a decoded command can benefit from shared control.

A central question raised by these results is whether the observed benefit could be reproduced by simpler forms of assistance. Explicit collision avoidance contributed substantially to the improvement in success, but an upper-bound analysis in which every BCI-only trial classified as blocked at an obstacle was reclassified as successful still did not reproduce shared-control performance, indicating that the benefit extended beyond the hard-coded safety mechanism. Similarly, fixed blending and phase-specific assistance could perform well under specific conditions but did not reproduce the performance profile of full shared control across environments, whereas prior-only control became increasingly unable to select the designated target as the number of candidate goals increased. Together, these analyses indicate that the shared-control benefit depended on continuous integration of the decoded BCI command with temporal and environmental information rather than on collision avoidance, a fixed amount of assistance, or autonomous trajectory generation alone.

Previous intracortical BCI studies in non-human primates have demonstrated complex continuous control under direct neural decoding, including three-dimensional cursor control and multi-jointed prosthetic-arm movements ^22,23^. Particularly relevant to the present study, Gilja et al. demonstrated direct two-dimensional BCI cursor control in a maze task requiring curved trajectories around visually defined barriers using ReFIT Kalman filter (ReFIT-KF^3^, and our previous work similarly demonstrated obstacle-avoiding virtual navigation without downstream shared assistance ^4^. Thus, shared assistance is not required to generate complex or obstacle-avoiding BCI trajectories. Rather, our results show that downstream shared control can improve the reliability with which continuously decoded commands are translated into successful actions under environmental constraints. This complements non-invasive BCI shared-control studies, in which autonomous navigation assistance has often been used to compensate for sparse or discrete neural commands ^6–8,10^. In our continuous intracortical setting, shared control strongly reduced execution-level failures but provided little benefit when the decoded command consistently favored a non-cued target, indicating that assistance acted primarily as an execution-stabilizing layer rather than correcting systematically misdirected decoded commands.

Previous invasive BCI shared-control studies have improved reaching and manipulation using sensor-based reflexes, vision-guided grasp assistance, or task-specific allocation of control between the user and robotic system ^15–17,19^. In our navigation paradigm, RT-V2 combined the decoded BCI velocity with a probabilistic trajectory prior derived from recent movement history and environmental context and was not provided with the identity of the cued target. Complete exclusion of the BCI command was restricted to the brief imminent-collision safety override; during standard shared control, the decoded BCI velocity continuously contributed to the posterior trajectory distribution.

Previous shared-control approaches often resolved uncertainty by shifting greater control authority to the assistive controller, either through fixed blending, phase-specific assistance or explicit goal inference ^13,15,16,19^. Such strategies can be effective when decoded commands are discrete, low-dimensional or unreliable, but they may be less appropriate for high-bandwidth invasive BCIs in which the decoded velocity carries meaningful moment-to-moment information about intended movement. In this setting, assistance should not determine the trajectory on behalf of the user, but should regulate how noisy decoded commands are expressed in the effector. The present framework implements this principle by continuously incorporating the decoded velocity while selectively suppressing execution-level instability induced by neural variability or environmental constraints. Thus, for high-bandwidth BCIs, shared control may be most effective when it stabilizes action execution under uncertainty rather than substituting user decisions.

The temporal integration used in our shared-control design shares a general principle with Kalman-filter-based BCI decoders such as ReFIT-KF, which use the previous state estimate to predict the current state before incorporating new neural observations^3^. Our PSID-based neural decoder likewise incorporates temporal dynamics through recursive estimation of behaviorally relevant latent states ^4,24^. RT-V2 therefore introduces an additional temporal dependence downstream of decoding. The mechanisms operate at different levels and over different quantities: decoder-level dynamics constrain the evolving latent-state estimate, whereas RT-V2 constructs a multimodal prior over future trajectories from the preceding 400 ms of decoded movement together with candidate goals and environmental context, which is then combined with the instantaneous decoded BCI velocity.

The prior-confidence index α reflects the consistency of the controller’s recent action history. When recent actions are consistent with task demands, α increases, reflecting a more concentrated temporal prior consistent with stronger temporal stabilization and reduced variability in execution. Abrupt deviations produce a transient dip in α, indicating increased uncertainty. This mechanism explains both the stabilization observed in obstacle-based tasks and the reduced responsiveness during goal changes: temporal priors improve control when recent action history remains predictive, but can impair responsiveness when that history becomes misleading.

Rapid goal changes reveal this limitation directly. In the Respawn task, the controller received no explicit signal of target reassignment and could update its posterior only in response to changes in the decoded velocity. Under these conditions, temporal-history dependence could contribute to delayed reorientation, but did not impose a fixed response delay: on successful trials, executed commands reoriented without detectable median additional delay relative to decoded BCI commands at 50 ms resolution. Failures were also associated with less reliable and slower reorientation of the decoded BCI command itself, including during BCI-only control. Offline prior-reset analyses nevertheless showed that removing pre-jump history improved shared-control responsiveness, demonstrating a specific contribution of the temporal prior. Thus, the limitation reflected contributions from both the incoming decoded command and the controller’s temporal history, rather than an unavoidable consequence of temporal stabilization.

These findings have direct implications for translating invasive BCIs from constrained laboratory tasks to real-world assistive devices. Outside the laboratory, users must control effectors over extended periods while neural signals, engagement and environmental demands fluctuate. In this setting, safe operation requires distinguishing transient variability in decoded commands from genuine changes in intent. The controller requires no additional neural information beyond the decoded command; its only extra input is environmental structure, which could be obtained from onboard cameras or depth sensors in assistive devices such as powered wheelchairs. By stabilizing execution without overriding the decoded command, shared control may reduce user burden and mitigate user–system conflict during prolonged use, particularly for devices such as powered wheelchairs, robotic manipulators and exoskeletons. Shared control therefore functions not as a substitute for neural control but as an enabling layer for reliable continuous control in complex environments.

Several steps remain before this approach can be deployed in clinical assistive devices. In structured environments such as a user’s home, static geometry and common destination locations could be mapped in advance, while existing computer-vision and depth-sensing methods could update obstacles and available goals online. RT-V2 could then operate on this continuously updated environmental representation, although perceptual uncertainty was not modeled here. RT-V2 supported variable goal configurations in simulation, although correct-target performance declined as the number of simultaneous candidate goals increased, particularly for larger candidate sets (10–20 targets) that extended beyond the candidate-set sizes represented during simulation pretraining. Generalization beyond this training distribution remains to be established. Because simulated pretraining requires some prior knowledge of the relevant task structure, more open-ended settings in which such structure is not known in advance would require additional methods. Moving obstacles, target occlusion, and perceptual errors were not explicitly evaluated here and remain important directions for future work^21^. Translation to people with paralysis will also require direct evaluation of perceived agency and subjective workload alongside task performance. Nevertheless, because the arbitration layer operates downstream of the neural decoder, the framework is not tied to a specific decoder or virtual effector. Any BCI system that provides a continuous command signal could, in principle, be coupled to similar task-specific priors to stabilize execution while preserving user control. In this way, confidence-modulated shared control provides a general post-decoder strategy for making continuous neuroprosthetic control safer and more reliable in dynamic environments.

## Methods

### Surgery, recording procedures and setup

Two adult male rhesus monkeys (*Macaca mulatta*, 7–9 kg) were implanted with a titanium headpost and three 96-channel Utah arrays (1–1.5 mm length, 400 µm spacing; Blackrock Neurotech), targeting F2 (PMd), F5c (PMv), and M1 (**Figure 1**a). Sex was not included as an experimental variable, and no sex-based analysis was performed because both animals were male and the study was not designed to assess sex-related differences. Surgeries were performed under propofol anesthesia (10 *mg kg*^−1^ ℎ^−1^) using stereotactic guidance, and verified via postoperative 3 T magnetic resonance imaging (MRI) (0.6 mm). Procedures were approved by the KU Leuven local ethical committee on animal experiment (approval number: P047/2021) and complied with NIH and EU animal care guidelines (Directive 2010/63/EU). Neural signals were acquired with Cereplex M headstages and a Cerebus system (256 channels, Blackrock Neurotech), filtered at 750 Hz, sampled at 30 kHz, and manually thresholded every session to extract spike activity.

Monkeys performed tasks while seated with head fixation, viewing a Viewpixx 3D display (1920×1080, 120 Hz) through synchronized shutter glasses for stereoscopic vision (Figure 1B). Full methodological details, including the virtual environment design and behavioral task structure, are available in Saussus et al.^4^.

### Task description

Building on our previous paradigm^4^, we introduced a shared-control mode to complement direct brain-controlled navigation. In this updated design, the BCI-decoded velocities were passed to a shared-control module, which adjusted movement commands in real time (**Figure 2**) (shared-control condition). The goal remained to navigate a sphere from a fixed starting point to a pseudorandomly placed target within a 3D environment.

### Experimental phases

During the **Passive fixation phase**, the monkey passively observed task execution performed by Unity’s AI navigation (NavMesh) system within a 3D virtual environment. To sample distinct movement directions, three targets (far left, straight, and far right) were used, with ∼30 trials per target collected while recording neural activity from three brain areas (M1, PMv, and PMd), as well as sphere velocity. In the **Decoder Training phase** (lasting up to one minute), a model was trained to predict movement velocities from the recorded neural signals. The subsequent **Online Decoding phase** included two conditions: (1) a BCI-only condition, in which the monkey directly controlled the movement of the sphere via real-time decoding of its neural activity, and (2) a shared-control condition in which the BCI-derived velocities were further processed by a shared-control module. This setup established a closed-loop BCI, with the monkey observing the resulting movements of the sphere in real-time on the 3D screen.

### Tasks

Each task was performed in a 3D environment and involved navigating a sphere along a two-dimensional (2D) plane towards the target represented by a white cube (Figure 1c). Within each recording session, all tasks shared a single Passive Fixation phase: during this phase, the sphere moved in a straight line from a fixed starting point toward one of three pseudorandomly presented targets (far left, straight, and far right), while the monkey passively observed the movement and maintained visual fixation. The dataset collected during this phase was used to train a single session-specific decoder, which was then applied—without retraining—across all subsequent tasks and conditions within that recording session.

During the Online Decoding phase, the monkey controlled the sphere using decoded neural activity (without any physical movement) to reach one of five pseudorandomly presented target positions. In the Fixed Obstacle task, a static obstacle (a black cube smaller than the target; Figure 1c) was placed between the starting point and the target, disrupting the straight-line path.

This task included interleaved trials of BCI-only and shared-control, recording 100 trials in total per session. In the Appearing Obstacle task (Figure 1D), the obstacle appeared in front of the sphere once it crossed an invisible threshold (z-coordinate = 1). At that moment, the obstacle was positioned at one-third of the distance from the sphere’s current position toward the cued target, such that its location depended on the sphere position at the time of appearance. To prevent anticipatory behavior, this task featured three pseudorandomly presented trial types: BCI-only without obstacle, BCI-only with obstacle and shared-control with obstacle, recording a total of 150 trials per session (i.e. 50 per condition; 10 per target across five targets). In all original Appearing Obstacle sessions in both monkeys (“Appearing Obstacle (December)”), shared-control trials additionally included a hidden fixed obstacle located halfway between the start position and the target that was considered by the controller but not rendered visible to the monkey. The obstacle that subsequently appeared during the trial was positioned separately based on the sphere’s current position and therefore did not coincide with this hidden fixed obstacle. This hidden fixed obstacle was absent in the follow-up experiment six months later in Monkey 1 (“Appearing Obstacle (6 months later)”). The Fixed and Appearing Obstacle tasks were performed sequentially within each session using the same decoder. To assess long-term effects of shared-control, the Appearing Obstacle task was repeated in Monkey 1 six months later using the same task structure, except that the hidden fixed obstacle was removed.

The Respawn task followed the same structure. After the sphere crossed the spatial threshold (z = 2, Figure 1e), the target could disappear and reappear (i.e. respawn) at an adjacent target location. This task also included three pseudorandomized conditions: no respawn, respawn with BCI-only control, and respawn with shared-control, recording a total of 210 trials per session (i.e. 70 trials per condition; 7 possible jump configurations, 10 trials each). The full experimental protocol has been described in detail in our previous study [Saussus et al.]^4^.

### BCI decoding algorithm

The decoding algorithm translated neural activity of PMv, PMd and M1 into real-time control of the sphere. The training of the decoding algorithm involved a supervised method, based on the Preferential Subspace Identification (PSID) algorithm developed by Sani et al. ^24^, using labeled data from the Passive Fixation phase. The data included 50 ms binned spike rates from electrodes —manually selected at session start based on the presence of spiking activity—paired with simultaneous sphere velocities *v* = (*v_x_*, *v_y_*, *v_z_*) and positions. Because the recorded extracellular signals and the set of active electrodes could vary across recording sessions, a brief Passive Fixation phase was performed at the beginning of each session and used to train a new session-specific decoder from the neural signals recorded that day. Once trained, the decoder produced continuous velocity updates every 50ms during online control using a previously validated non-linear, real-time extension of PSID, and its parameters were held fixed across all subsequent tasks and conditions within that recording session. During online decoding, the behaviorally relevant PSID latent state was estimated recursively using a Kalman-filter update: the previous latent-state estimate was propagated through the learned dynamics to predict the current state, which was then updated using the current neural observation before being mapped to velocity. Full implementation details are provided in Saussus et al.^4^.

### Shared-control paradigm

RT-V2^21^ was implemented as a post-processor to the neural BCI decoder and operated at 20 Hz during closed-loop control.

The intent predictor modeled a prior distribution *p*(*ξ*|ℎ*_t_*, *c*), which represents candidate future velocity trajectories *ξ* conditioned on the environmental context *c* and recent movementhistory ℎ*_t_* (last eight 50 ms steps, 400 ms total). More specifically, RT-V2 represented the velocity action distribution at each future time step as a Gaussian mixture,

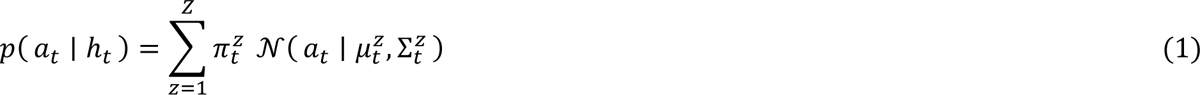

where *z* indexes maneuver modes and π^z^_t_, μ^z^_t_, and Σ^z^_t_ denote the corresponding mixture weight, mean velocity, and covariance. Sampling these velocity distributions over the prediction horizon generated candidate future trajectories *ξ*. The trajectory model was pretrained on the Trajectory-1M simulation dataset described by Song et al.^21^, comprising 1,000,000 successful navigation trajectories (900,000 training; 100,000 validation). Training environments contained 1–6 candidate targets and 1-6 static obstacles. Trajectories were generated in randomized target and obstacle configurations using Probabilistic Roadmap paths tracked by a noisy proportional controller. Target and obstacle locations were sampled independently, such that obstacle location did not provide a systematic cue to the designated target.Trajectory-1M was partitioned using a fixed random seed, with scene-disjoint assignment across data splits such that evaluation environments were not observed during training. For the present online BCI experiments, the pretrained model was transferred without fine-tuning on monkey BCI trajectories.

The model learned to predict plausible future trajectories given environmental context and past motion history. Neural activity was not incorporated during prior training; only decoded kinematics and task context were used. The prior was conditioned on the set of five candidate targets but did not encode which target was cued. At trial onset, the prior supported trajectories toward all candidate targets; as motion history accumulated, the distribution progressively concentrated on trajectories consistent with recent velocity patterns and environmental feasibility. Importantly, the instantaneous decoded velocity *v* was not incorporated in computing the prior.

During online control (**Figure 2**), the system first generated the prior *p*(*ξ*|ℎ*_t_*, *c*), from environmental context and motion history. The decoded velocity *v* was then incorporated through a likelihood model *p(v|ξ;Σ_sys_)*, which quantified how compatible each candidate trajectory was with the observed command. At each time step, a trajectory *ξ* implies a specific instantaneous velocity; trajectories whose implied velocity closely aligned with the decoded velocity were assigned higher likelihood, whereas trajectories that diverged from it received lower probability. The decoded BCI velocity was modeled as a Gaussian observation of the instantaneous velocity implied by the candidate trajectory:

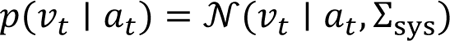

where *a_t_*is the instantaneous velocity implied by the candidate trajectory and Σ_sys_specifies the assumed uncertainty in the decoded BCI command. The posterior distribution was computed via Bayes’ rule,

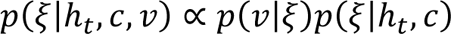

Candidate trajectories were sampled from this posterior and evaluated using a cost function that penalized obstacle collisions and rewarded progress toward candidate targets. The trajectory minimizing cost was selected, and its instantaneous velocity was executed.

Prior concentration was summarized using the confidence index *α_t_*, computed from the entropy lower bound *H^LB^_t_* of the prior action distribution. For each trial independently, *H*_max_ was estimated from the entropy samples included in the corresponding analysis:

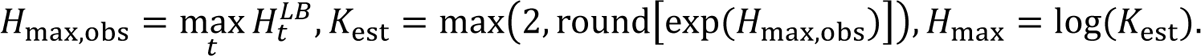

The confidence index was then defined as

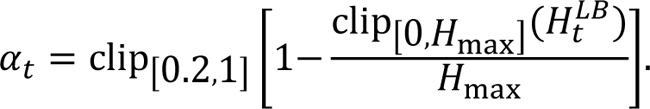

If the maximum observed entropy within a trial was invalid or non-positive, the 95th percentile of that trial’s observed entropy values was used instead to estimate *K*_est_. Thus, *α_t_* = 1 corresponds to a highly concentrated prior, whereas the imposed lower bound *α_t_* = 0.2 represents high-entropy prior states. The confidence index *α_t_* was used as a descriptive measure of prior concentration and was not itself an input to the controller or an explicit blending coefficient.

The relative contribution of the prior and decoded BCI command to the posterior depended on their respective distributions and on the system covariance parameter Σ_sys_. For a fixed Σ_sys_, a more concentrated prior (low entropy) can exert greater influence on the posterior, but prior concentration alone does not uniquely determine the resulting change in executed velocity. Identical *α_t_* values can therefore produce different executed-velocity corrections depending on the instantaneous BCI command and its relationship to the prior. Conversely, a broader, high-entropy prior reflects greater uncertainty in the predicted action distribution and can allow the instantaneous decoded BCI command to exert greater influence on the posterior. Σ_sys_ specified the assumed uncertainty in the decoded BCI velocity: higher values reduced the influence of the BCI likelihood relative to the prior, whereas lower values had the opposite effect. The system covariance parameter Σ_sys_ was selected empirically during pilot testing with Monkey 1. During pilot sessions, we tested multiple candidate values and selected the setting that provided stable shared-control behavior while retaining responsiveness to the decoded BCI command. We used:

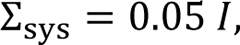

where *I* is the 2×2 identity matrix. This value was then held fixed across all subsequent sessions and tasks in Monkey 1 and was used unchanged for Monkey 2.

In safety-critical situations—defined as cases in which the decoded velocity would lead to an imminent collision with an obstacle— the arbitration mechanism briefly shifted fully toward the shared-control prior, resulting in a brief safety override. During this override, which lasted for three consecutive update steps (150 ms total), the executed velocity was determined exclusively by the shared-control prior, without incorporating the current BCI command. The controller then returned immediately to the standard shared-control regime.

For the invasive BCI implementation, the continuously decoded two-dimensional BCI velocity served as RT-V2’s user-command input. Apart from this input and the imminent-collision safeguard described above, the underlying probabilistic prior, likelihood, posterior inference, and trajectory-selection architecture were otherwise unchanged.

### Simulation and controller-comparison analysis

#### Fixed-blending and prior-only comparison

To compare full shared control with alternative control strategies, we performed closed-loop simulations at 10 Hz. At control step *t*, the simulated user generated a velocity command directed toward the designated target *g*:

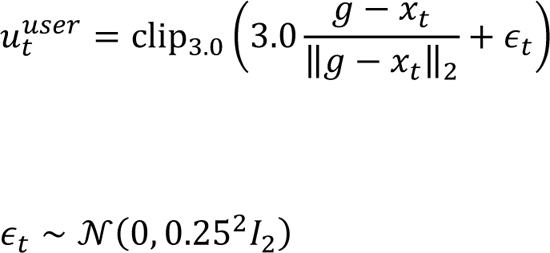

where *x_t_* is the current sphere position and *clip*_3.0_ rescales commands with norm greater than 3.0 *ms*^−1^ to a norm of 3.0 m *ms*^−1^. When the sphere coincided with the target, the nominal target-directed component was set to zero.

For fixed-blend control, the executed command was

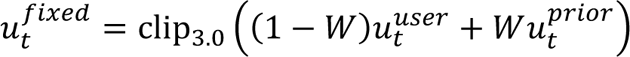

where *u^t^_prior_* was the action generated from the trajectory prior without conditioning on the instantaneous simulated-user command. We evaluated fixed prior weights *W* = 0.25, 0.50, 0.75, 0.80, 0.85, 0.90 and 0.95, together with user-only, prior-only, and full shared-control conditions. Before the trajectory model accumulated the four positions required for its first prediction, the adapter returned the incoming user command; the fixed-blend output during these warm-up samples therefore reduced to the clipped user command. In the user-only condition, *u^t^_user_* was applied directly; in the prior-only condition, actions were generated without conditioning on the instantaneous user command; and full shared control used the standard probabilistic posterior controller. The imminent-collision safety override was retained for full shared control but disabled for the fixed-blending baseline.

Simulations were evaluated in two environments. In the BCI-like environment, five candidate target locations were positioned at (6,-7), (8.5,-3.5), (9.2,0), (8.5,3.5) and(6,7)*m*. Target identities were permuted across trials, with one target designated as the true target, and a single static obstacle was positioned halfway between the starting position and the true target. The randomized cluttered simulation environment comprised a 13 *x* 18*m* workspace with the sphere initialized at the origin. Between 1 and 6 cubic obstacles (side length 1.3 m) were positioned between radial distances of 3 and 6m from the origin, whereas 1–6 candidate targets were positioned beyond 6m and within the workspace boundaries.

One hundred matched trials were evaluated per controller condition in each environment. Scene configuration, designated target, simulated-user noise realization, and random seed were held identical across controller conditions. Sphere position was updated as *x_t_*_+1_ = *x_t_* + 0.1*u_t_*. Trials terminated after a maximum of 200 control steps (20 s), upon collision, or upon target acquisition. Collision was defined as planar distance < 1.3*m* from any obstacle. Success was defined as planar distance < 1*m* from the designated target. Collision was evaluated before target acquisition when both criteria were met after the same update; trials satisfying neither criterion within 200 steps were classified as timeouts. The same pretrained trajectory model and controller settings were used across all controller conditions; only the manner in which the simulated-user command was incorporated differed between conditions. Paired success-rate differences and 95% bootstrap confidence intervals were computed by resampling matched trial pairs, and binary success and collision outcomes were compared using two-sided exact McNemar tests.

#### Hard-coded phase-specific assistance

Using the same matched simulation framework as the fixed-blending comparison, we evaluated a spatially gated controller in which the simulated user command was executed when the distance to the nearest obstacle was at least *R*, and the RT-V2 prior action was executed when this distance fell below *R*. We tested *R* = 1.5, 2.0, 2.5 and 3 *m*. All other simulation settings and outcome definitions were identical to the fixed-blending comparison.

#### Candidate-target scalability and prior-only control

To assess how target ambiguity affected performance, we compared full shared control with prior-only control while varying the number of candidate targets (1, 2, 3, 4, 5, 10, 15, and 20; 100 trials per candidate-set size and controller condition). Targets were sampled uniformly within [0,13] *x* [−9,9] *m* and constrained to lie >6 m from the start and ≥1 m from all other targets; the first target was designated as the true target. No obstacles were present. The same target configurations and trial seeds were used for both controller conditions. Prior-only control retained candidate-target information and position history but was not conditioned on the instantaneous user command. Trials terminated when the sphere came within 1.0 m of any candidate target or after 200 control steps. Correct-target success denoted acquisition of the designated target, whereas any-target success included acquisition of either the designated target or a distractor.

#### Robustness to user-command reliability

To assess robustness to directional variability in the user command, we used the randomized cluttered simulation environment described above, containing 1–6 candidate targets and 1–6 static obstacles per trial. The simulated-user command had a fixed magnitude of 3*ms*^−1^ toward the designated target, with its direction perturbed independently at each control step by a zero-mean Gaussian angular error with standard deviation 0°, 10°, 20°, 40°, or 60°. User-only and full shared-control conditions were evaluated using 100 trials per angular-error level, with the same scenes and matched random realizations across conditions. Success was defined using the same 1-m target-acquisition criterion as above.

#### Time-resolved influence of the user command

To assess whether the instantaneous user command continued to influence shared control throughout movement, we performed 100 simulations using the BCI-like five-target geometry described above, with a single obstacle positioned on the ray from the start to the designated target. Success was defined using a 1.0 m target-acquisition criterion, and the standard imminent-collision safeguard was retained. The simulated user generated a deterministic 3 *m s*^−1^ command toward the designated target at each control step. Movement phase was defined from projected progress along the start-to-target axis as early (<33%), middle (33–67%), or late (≥67%). During prior-only rollouts, the prior action and the user-command-conditioned posterior action were evaluated at each visited state while holding the controller and random state fixed; the environment was advanced using the prior action only. User-command influence was quantified both as the magnitude of the posterior–prior action difference and as its cosine similarity with the instantaneous user command. Alignment was averaged within each trial and phase, and trial-level means were tested against zero using two-sided one-sample t-tests.

In a complementary late-command perturbation, 100 matched posterior-controlled trial pairs were compared. In baseline trials, the 3 *m s*^−1^ command remained directed toward the designated target throughout movement; in perturbed trials, it was replaced after 67% projected progress by a sustained 1 *m s*^−1^ command in the fixed direction *d* = (0, −1). Matched trials used identical scenes, controller seeds, and pre-perturbation commands. Baseline and perturbed trajectories were compared over their shared post-perturbation duration. Trajectory divergence was quantified as the mean per-step Euclidean distance between the matched baseline and perturbed positions (average displacement error, ADE), and tested against zero using a two-sided one-sample t-test. For each trajectory, cosine similarity between consecutive movement vectors and the imposed direction was averaged within trial, and baseline and perturbed trial-level values were compared using a two-sided paired t-test. Ninety-five percent confidence intervals for trial-level means were calculated using the t distribution.

### Analysis

#### Success rate and learning trends

For each session and condition (shared-control, BCI-only), success rate was defined as the fraction of trials in which the sphere dwelled in the cued target window for 500 ms. Session-wise success rates were compared between shared-control and BCI-only conditions using paired, two-sided Wilcoxon signed-rank tests across sessions (n = 9–12 per task). To estimate chance performance, we conducted a permutation test (10,000 permutations; test statistic: success rate across trials). Because each target was always paired with a specific obstacle in the Fixed and Appearing Obstacle tasks, permutations were performed over joint (target, obstacle) pairs. Learning trends were assessed using Spearman’s rank correlation between session index and success rate, computed separately for shared-control and BCI-only conditions.

#### Per-target shared-control gain and scaling behavior

For each session and target, success rates were computed separately for shared-control and BCI-only trials. Per-target shared-control gain (Δ) was defined as the difference between shared-control and BCI-only success rates and computed independently for each session. To assess how shared-control gain depended on baseline BCI performance, individual target × session observations from the Fixed Obstacle and Appearing Obstacle datasets, including the six-month follow-up, were analyzed using quadratic regression.

To assess whether the relationship between baseline BCI performance and shared-control gain was consistent across subjects and tasks, we additionally analyzed target × session observations while retaining monkey, task, and session identity. Quadratic regression models included linear and quadratic terms for baseline BCI success. Standard errors were clustered by session to account for multiple target observations within each recording session. We compared a common-shape model containing monkey and task intercepts with models allowing the linear and quadratic baseline terms to vary by monkey or task using likelihood-ratio tests, Akaike information criterion (AIC), Bayesian information criterion (BIC), and grouped cross-validation. Separate quadratic fits were also computed for each monkey and monkey × task dataset. The distribution of observations across baseline-performance ranges was quantified in 20-percentage-point bins.

To assess whether very low baseline task success reflected an absence of directional information in the decoded command, we performed a post hoc analysis of BCI-only trials from target × session observations with exactly 0% and >0–10% baseline success. For each BCI-only trial, directional alignment was quantified as the mean cosine similarity between the decoded velocity vector and the direction from the current sphere position to the cued target across time points with available decoded velocity. Samples preceding target onset were excluded when timing information was available. Trial-level cosine similarities were then summarized within each baseline-success group. We additionally quantified the number of BCI-only trials contributing to each target × session observation to characterize the sampling resolution of the baseline success estimate.

#### Trajectory-level performance metrics

To quantify how shared-control affected movement execution, we compared trajectory-level metrics between shared-control and BCI-only trials. Metrics included first target entry, total acquisition time, trajectory smoothness, obstacle clearance, and collision incidence. Time to first target entry was defined as the interval between movement onset and the first sample within the target window and was computed for all included trials that reached the target window. Total acquisition time was defined as the interval between movement onset and successful completion of the trial of the required 500-ms target hold, and was computed for successful trials only. Trajectory smoothness was quantified using mean squared jerk (MSJ), computed from position trajectories and *log*_10_-transformed prior to analysis. Obstacle clearance was defined as the minimum sphere–obstacle distance over the trajectory, and collisions were defined as any time point at which the sphere intersected the obstacle. Path-length efficiency was defined as the minimum straight-line distance from movement onset to the edge of the target window divided by the executed path length. Total acquisition time was measured from movement onset until successful completion of the 500 ms target hold, and mean movement speed was computed over the executed trajectory for successful trials000. Metrics were first averaged within session and condition, and session-level differences between shared-control and BCI-only trials were assessed using paired, two-sided Wilcoxon signed-rank tests. For visualization, trajectories were temporally aligned, interpolated to a common time base, averaged per target, and grouped by avoidance side.

#### Failure modes and decoder-inferred choice

To characterize how shared-control affected unsuccessful behavior, we performed a post hoc classification of all unsuccessful trials, defined as trials in which the monkey failed to dwell in the cued target window for at least 500 ms. We distinguished behavioral outcomes, which describe how the executed trajectory terminated, from decoder-inferred choice states, which summarize the directional content of the decoded BCI command across the trial. Each unsuccessful trial was assigned to a single, mutually exclusive behavioral failure category based on trajectory geometry and task-defined outcomes. Trials were classified as “blocked at obstacle” if the trajectory entered an inflated obstacle region (obstacle half-side plus avatar radius and margin) for at least ten consecutive frames without passing the obstacle in the forward direction. Among the remaining trials, “target overshoot” was assigned if the trajectory entered the cued target window but subsequently exceeded its far edge along the forward (+z) direction. Trials were labeled “insufficient dwell time” if the cued target window was entered but the longest continuous dwell was shorter than 500 ms. Trials were labeled “non-cued target acquisition” if they achieved a continuous dwell of at least 500 ms within the target window corresponding to a non-cued candidate target location, without overlapping the cued target window. Trials were labeled “insufficient target proximity” if the final position was closest to the cued target among all targets, and the maximal forward position remained below the near edge of the cued target band. Any remaining unsuccessful trials were grouped as “other”.

To relate trial outcomes to the quality of the decoded command, we assigned each trial a decoder-inferred choice state based on the decoded BCI velocity. During online control, the monkey viewed only the cued target; the other candidate target locations were available to the shared-control controller but were not displayed to the monkey. At each time step, the decoded velocity vector was compared to vectors pointing from the current position to each candidate target. For the obstacle tasks; time steps for which the sphere center was within 1.5 Unity units of the obstacle center in the x–z plane were excluded from this analysis, and for the Respawn task only post-jump time steps were included. The trial-wise choice distribution was then defined as the fraction of included time steps for which each target direction yielded the maximal positive cosine similarity. Trials were labeled “correct choice” when the cued target accounted for ≥60% of included time steps; “incorrect choice” when the direction of a specific non-cued candidate target accounted for ≥60%; and “ambiguous choice” otherwise. Ambiguous trials were further labeled “adjacent choice” when ≥80% of the non-true choice mass was concentrated on the immediate geometric neighbors of the cued target, and “ambiguous choice” otherwise. Decoder-direction error refers more broadly to trials in which the decoded BCI velocity was not consistently aligned with the cued target direction and therefore includes adjacent, ambiguous, and incorrect choice states. These choice-state labels summarize the trial-wise distribution of candidate target directions favored by the decoded BCI velocity and are distinct from the final behavioral outcome; for example, an “incorrect choice” indicates that the decoded command predominantly favored the direction of a specific non-cued candidate target but does not imply that this location was ultimately acquired. We then computed success probability separately for shared-control and BCI-only trials within each choice state, yielding *P*( *success* ∣ *c*ℎ*oise state*, *condition*).

To estimate the maximum contribution of collision prevention alone, we performed an upper-bound counterfactual analysis on the empirical BCI-only trials. Every trial classified as an “blocked at obstacle” failure was reclassified as successful, while all other trial outcomes were left unchanged. The resulting counterfactual success rate was compared with the observed shared-control success rate; the residual difference represents the portion of the shared-control gain that could not be explained by perfect rescue of “blocked at obstacle” failures alone.

#### Event-locked confidence and rapid goal-change analysis

To characterize how prior confidence responded to abrupt environmental changes, we analyzed the temporal evolution of the shared-control prior confidence index α around task-defined events. At each time point, *α*(*t*) was defined as one minus the normalized entropy lower bound of the shared-control prior’s action distribution. For the Appearing Obstacle task, α was aligned to obstacle onset; for the Respawn task, *α* was aligned to target respawn (t = 0).

For each trial, a pre-event baseline *α_pre_* was computed as the mean α over the −400 to 0 ms window. Within the 0–800 ms post-event window, we identified the minimum α value (*α_min_*) and defined the dip magnitude as Δ*α_dip_* = *α_min_* − *α_pre_*. Recovery time was defined as the first post-minimum time point at which *α*(*t*) exceeded *α_pre_*; trials without recovery within the analysis window were excluded from recovery analyses. To control for slow within-trial fluctuations unrelated to the event, we repeated the same analysis using pseudo-events placed at random, non-overlapping times within each trial and computed Δ*α_pseudo_* = *α_post_* − *α_pre_*, where *α_post_* was the mean α in the corresponding post-event window, from the corresponding baseline and post-event window. Statistical comparisons were performed on session-level summary metrics using paired, two-sided Wilcoxon signed-rank tests. Shaded confidence intervals shown in **Figure 6** and **Extended Data Figure 4** reflect across-trial variability for visualization only; statistical inference was performed on session-level summary metrics using paired, two-sided Wilcoxon signed-rank tests. Confidence intervals for the across-trial mean were calculated as mean ± 1.96 SEM.

To examine how temporal arbitration responds to abrupt goal changes, we analyzed the Respawn task in which the cued target abruptly changed during the trial. Reorientation was quantified separately for the decoded BCI velocity and the executed velocity. At each post-respawn time point, directional evidence for the new relative to the previous target was computed as the difference between the velocity projection toward the new-target direction and the velocity projection toward the old-target direction. Reorientation latency was defined as the first post-respawn time point at which this evidence became positive and remained positive for at least 200 ms. A signal was classified as having reoriented within a given analysis window only when the full 200 ms sustained-evidence period occurred within that window. For the primary analysis, a 2 s post-respawn window was used.

To assess whether updating of the decoded BCI command was associated with behavioral outcome, we compared the proportion of trials in which the decoded BCI velocity reoriented and, among trials that reoriented, the reorientation latency between successful and unsuccessful trials separately for shared-control and BCI-only conditions. Robustness was assessed by repeating the classification using post-respawn windows from 1 to 3 s while keeping the 200 ms sustained-evidence criterion fixed.

To quantify additional delay between the decoded and executed commands, we analyzed shared-control trials in which both signals reoriented. For each trial, the BCI-to-executed reorientation lag was defined as *t_exec_* − *t_BCI_*, where *t_exec_* and *t_BCI_* denote the corresponding reorientation latencies. Positive values therefore indicated that reorientation of the executed command followed reorientation of the decoded BCI command.

To quantify persistence of the previous temporal prior, we computed a goal mismatch index (*GMI* = cos *θ_old_* − cos *θ_nenn_*), defined as the difference between the cosine similarity of the executed velocity to the old and new target directions. The decay time was defined as the first time point at which GMI ≤ 0, indicating alignment with the new goal. Decay times were compared between successful and unsuccessful shared-control trials using paired Wilcoxon signed-rank tests on session medians, with Benjamini–Hochberg false-discovery-rate correction across the tested Respawn metrics.

To isolate the contribution of the temporal prior, we performed offline replay of the same shared-control trials using identical neural data and decoded velocities, but with the temporal prior either carried over across respawn or reset at the moment of target jump. Reorientation lag and behavioral performance were compared between the two replay conditions.

## Data availability

All processed data supporting the findings of this study are available at https://doi.org/10.48804/7KGSQS25 and will be made public at the time of publication.

## Code availability

Analysis code is available on GitHub (https://github.com/ophelie-bci/Shared_control_analysis) and archived at Zenodo (doi: https://doi.org/10.5281/zenodo.1892613526). The RT-V2 shared-control framework used in this study was developed by Song et al.^21^. Source code for RT-V2 is publicly available at https://github.com/mousecpn/Robot-TrajectronV2, with the shared-control implementation, pretrained model and simulation analyses available at https://github.com/mousecpn/RTV2_ws.

## Acknowledgements

We thank S. Verstraeten, M. De Paep, W. Depuydt, I. Puttemans, and C. Ulens for technical assistance. We thank A. Hermans and S. De Pril for administrative support. This work was supported by Fonds Wetenschappelijk Onderzoek (FWO) grant G.097422N and KU Leuven grants C14/18/100 and C14/22/134. The text of the article was partially edited by a large language model (OpenAI ChatGPT).

## Author Contributions

Conceptualization: O.S., P.S., S.D.S., R.D., and P.J. Methodology: O.S., P.S., S.D.S., R.D., T.D., and P.J. Investigation: O.S., S.D.S., I.C., and P.S. Visualization, validation, and formal analysis: O.S. and P.S. Supervision: R.D., T.D. and P.J. Writing—original draft: O.S. Writing— review and editing: O.S., P.S., S.D.S., R.D., T.D., and P.J. Software: O.S. and P.S. Data curation: T.D., O.S., and S.D.S. Resources and funding acquisition: R.D., and P.J. Project administration: P.J. and S.D.S.

## Extended data

**Extended Data Figure 1.**
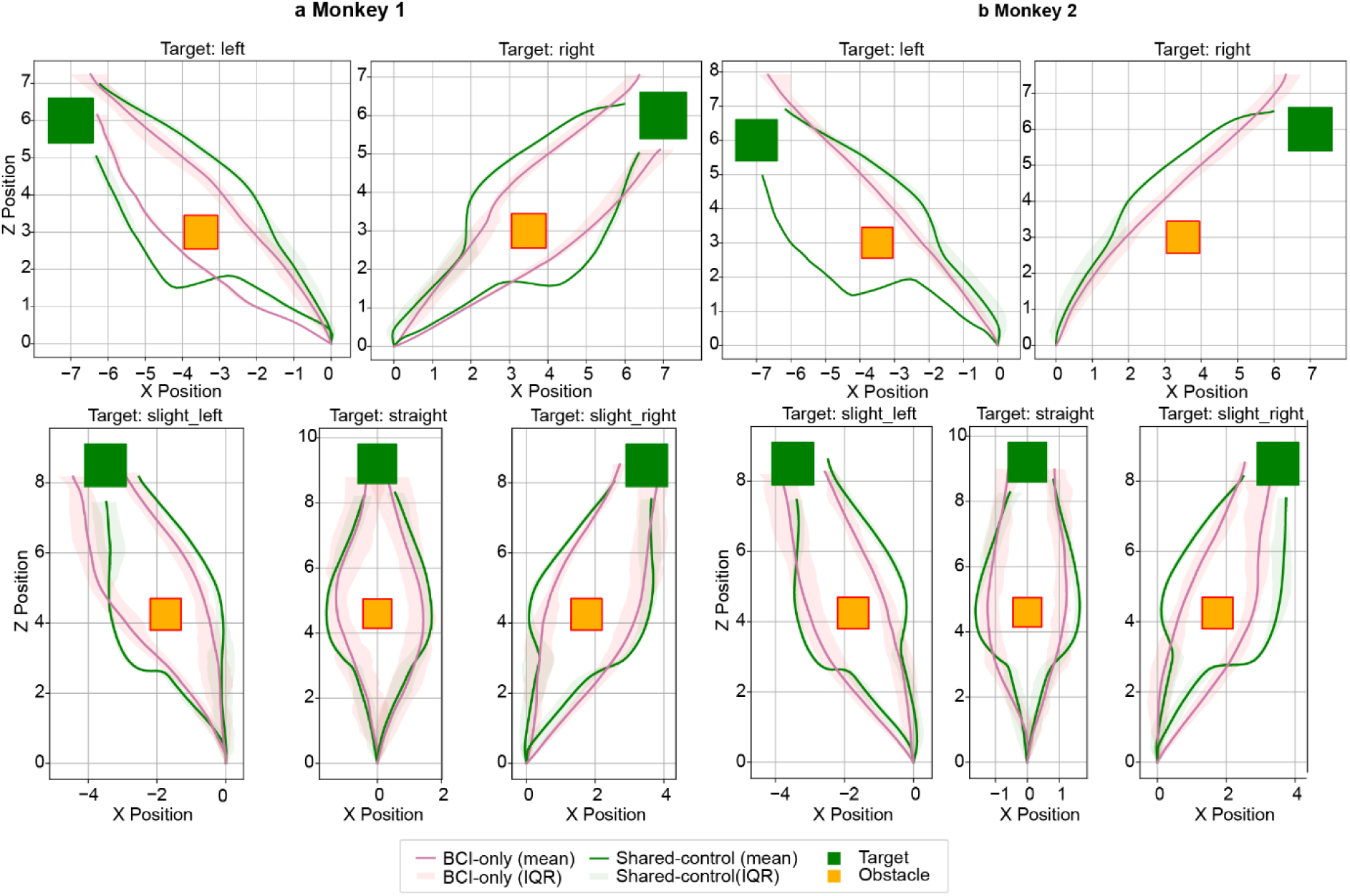
Shared-control alters obstacle-avoidance trajectories. **a, b**, Average trajectories of Fixed Obstacle task, shown separately for Monkey 1 (a) and Monkey 2 (b) across different target directions. Green traces indicate shared-control trials and pink traces indicate brain-computer interface (BCI)-only trials. Solid lines represent across-trial mean trajectories averaged over all sessions, and shaded regions denote interquartile ranges. Cued target locations (green squares) and obstacle positions (orange squares) are shown for reference. Across target configurations, shared-control produces smoother and more consistent trajectories that maintain larger clearance from obstacles, reducing overshoot and collision-prone deviations relative to unassisted control. Quantitative trajectory-level comparisons are reported in Extended data Table 1.

**Extended data Figure 2.**
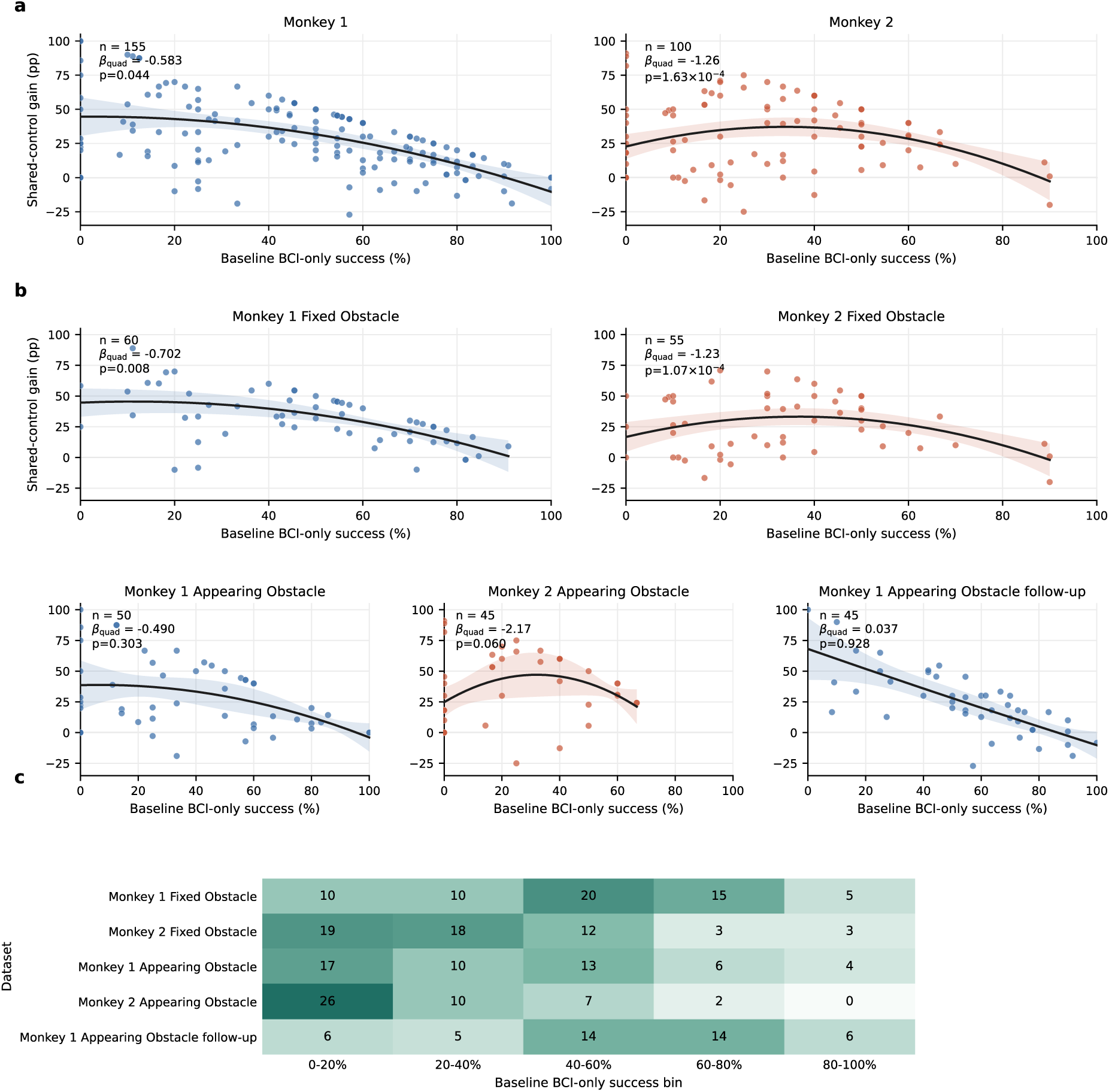
Dependence of shared-control gain on baseline brain-computer interface (BCI) performance across monkeys and tasks. **a**, Relationship between baseline BCI-only success and shared-control gain analyzed separately for each monkey. Each point represents one target in one session. Black curves show quadratic fits restricted to the range of baseline performance observed within each dataset, and shaded regions indicate S5% confidence intervals computed from the session-clustered covariance matrix. Values indicate the number of target × session observations (n), the quadratic coefficient (β_quad_), and the corresponding p-value, with standard errors clustered by session. **b,** Corresponding analyses performed separately for each monkey × task dataset. Axes are held constant across panels to illustrate differences in the sampled ranges of baseline performance. **c,** Number of target × session observations contributing to successive ranges of baseline BCI-only performance for each dataset. The complete analysis comprised 255 target × session observations from 51 sessions. Shared-control gain was defined as shared-control success minus BCI-only success. pp, percentage points.

**Extended data Figure 3.**
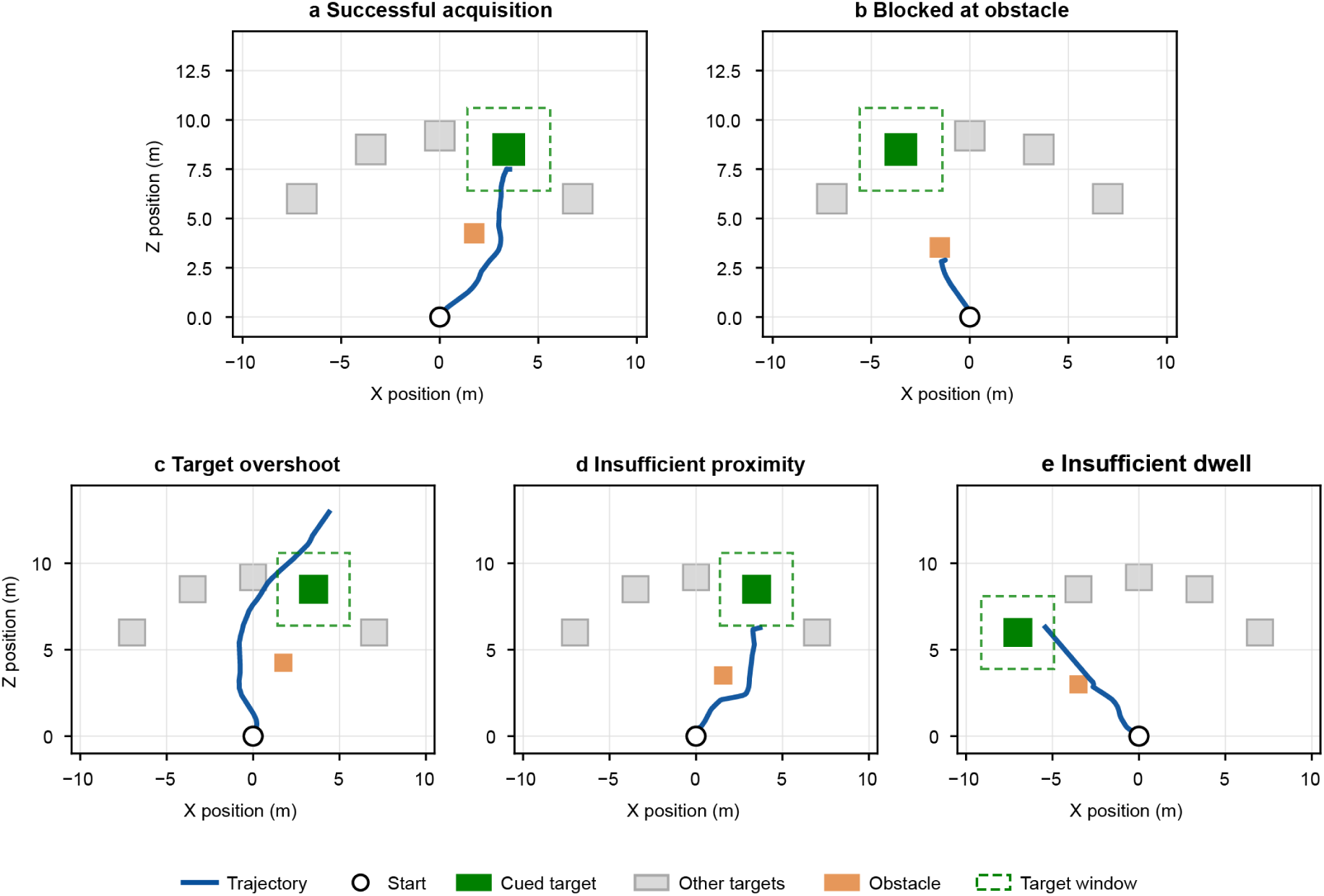
Representative examples of execution-related trial outcomes. Representative X–Z plane trajectories illustrating successful target acquisition and the execution-related failure categories quantified in Figure 5. **a,** Successful acquisition, in which the trajectory enters the cued target window and satisfies the required 500 ms dwell. **b,** Blocked at obstacle, in which forward progress is prevented by the obstacle. **c,** Target overshoot, in which the trajectory enters the cued target window but continues beyond it without satisfying the dwell criterion. **d,** Insufficient proximity, in which the trajectory approaches the cued target but does not enter the target window. **e,** Insufficient dwell, in which the trajectory enters the cued target window but remains there for less than 500 ms. Blue lines show executed trajectories and open circles indicate movement onset. The cued target is shown as a green square, other candidate targets as grey squares, obstacles as orange squares, and the cued target window as a dashed green outline.

**Extended Data Figure 4.**
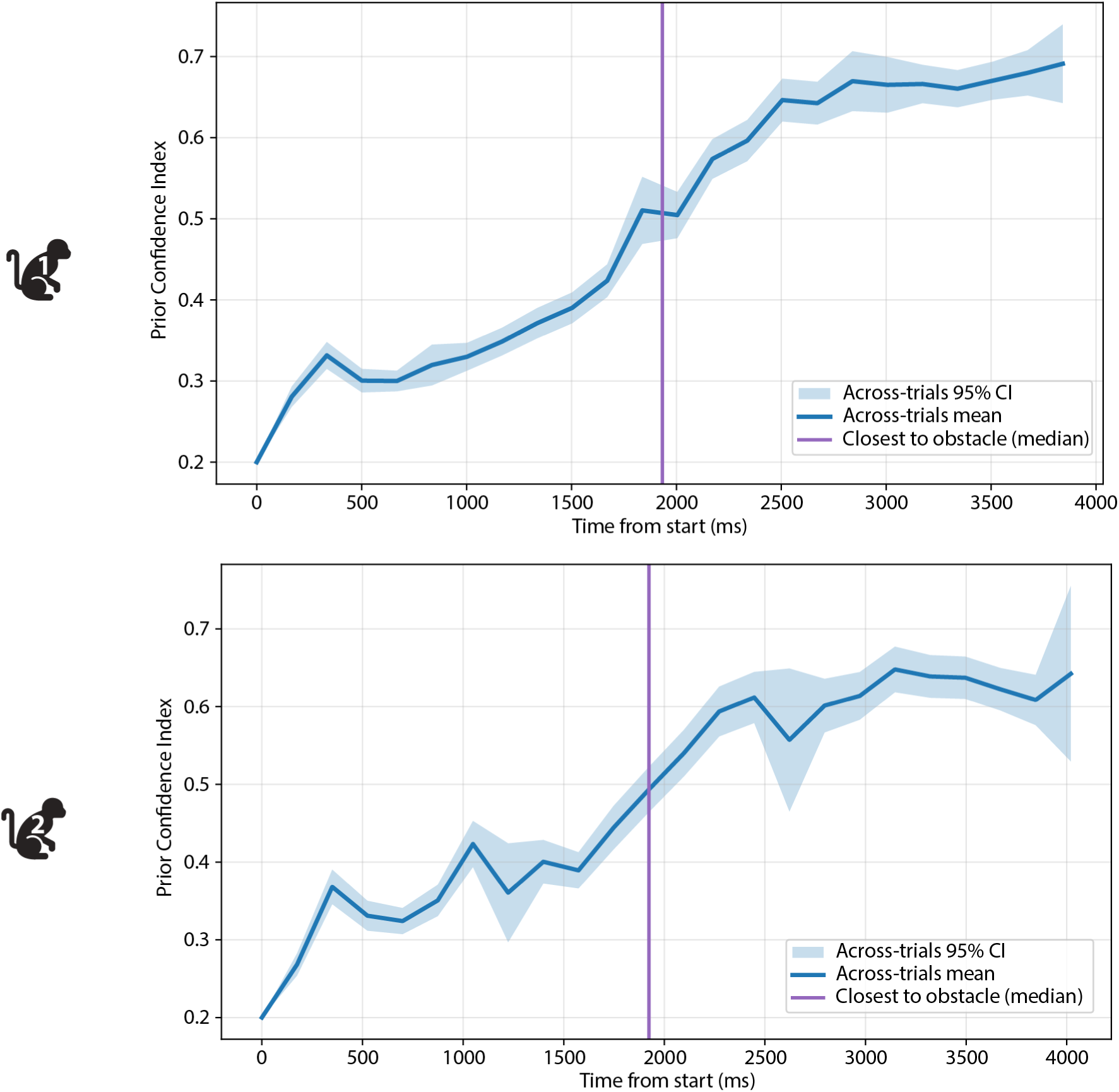
Absence of event-locked modulation of the shared-control temporal prior in the Fixed Obstacle task. Across-trial mean shared-control prior confidence index α (defined as 1 − normalized entropy of the prior action distribution) aligned to trial start for two subjects (top: Monkey 1; bottom: Monkey 2). Solid blue traces denote the across-trial mean, with shaded regions indicating the S5% confidence interval across trials. The vertical purple line marks the median time of closest approach to the obstacle across trials. In contrast to the Appearing Obstacle and Respawn tasks (**Figure Ca,b**), no transient reduction in α is observed around this time. Instead, α increases throughout the trial, consistent with stable prior confidence when environmental constraints are present from trial onset and recent movement history remains predictive. The number of sessions (N) for each task is provided in Extended data Table 2.

---

**Extended Data Table 1.** Trajectory-level effects of shared-control. Values are session-level paired differences (Δ = shared-control − BCI-only). Path efficiency, total acquisition time, and mean squared jerk were computed on successful trials; time to first target entry was computed for all trials that entered the target window; clearance and collision rate were evaluated separately for successful and unsuccessful trials. Collision-rate differences are in percentage points. P values are from paired, two-sided Wilcoxon signed-rank tests. N indicates the number of paired sessions.

| Dataset | Metric | N | Mean $\Delta$ | P |
| --- | --- | --- | --- | --- |
| Appearing Obstacle | Path efficiency | 19 | +0.0079 | 0.490 |
| | Time to first target entry (s) | 19 | +0.515 | $7.25 \times 10^{-5}$ |
| | Total acquisition time (s) | 19 | +0.682 | $1.14 \times 10^{-5}$ |
| | Speed (m/s) | 19 | -0.351 | $3.81 \times 10^{-6}$ |
| | Mean squared jerk | 19 | -149,843 | $2.67 \times 10^{-4}$ |
| | Clearance, successful (m) | 19 | +0.444 | $3.81 \times 10^{-6}$ |
| | Clearance, unsuccessful (m) | 18 | +0.551 | $7.63 \times 10^{-6}$ |
| | Collision rate, successful (pp) | 19 | -59.16 | $1.93 \times 10^{-4}$ |
| | Collision rate, unsuccessful (pp) | 18 | -70.50 | $7.63 \times 10^{-6}$ |
| Fixed Obstacle | Path efficiency | 23 | +0.0100 | 0.0605 |
| | Time to first target entry (s) | 23 | +0.473 | $2.38 \times 10^{-7}$ |
| | Total acquisition time (s) | 23 | +0.583 | $2.38 \times 10^{-7}$ |
| | Speed (m/s) | 23 | -0.329 | $2.38 \times 10^{-7}$ |
| | Mean squared jerk | 23 | -151,229 | $1.67 \times 10^{-5}$ |
| | Clearance, successful (m) | 23 | +0.481 | $2.38 \times 10^{-7}$ |
| | Clearance, unsuccessful (m) | 23 | +0.770 | $2.38 \times 10^{-7}$ |
| | Collision rate, successful (pp) | 23 | -48.85 | $2.38 \times 10^{-7}$ |
| | Collision rate, unsuccessful (pp) | 23 | -57.63 | $2.38 \times 10^{-7}$ |
| Appearing Obstacle<br>6-month follow-up | Path efficiency | 9 | +0.0235 | 0.0117 |
|  | Time to first target entry (s) | 9 | +0.687 | 0.00391 |
|  | Total acquisition time (s) | 9 | +0.850 | 0.00391 |
|  | Speed (m/s) | 9 | -0.461 | 0.00391 |
|  | Mean squared jerk | 9 | -212,612 | 0.00391 |
|  | Clearance, successful (m) | 9 | +0.479 | 0.00391 |
|  | Clearance, unsuccessful (m) | 9 | +0.669 | 0.00391 |
|  | Collision rate, successful (pp) | 9 | -66.18 | 0.00391 |
|  | Collision rate, unsuccessful (pp) | 9 | -74.66 | 0.00391 |

**Extended Data Table 2.** Task performance with and without shared-control assistance. Summary of success rates across tasks, sessions, and monkeys. For each task, the number of sessions is listed along with the mean success rate ± standard deviation (SD) for brain-computer interface (BCI)-only trials and shared-control trials. Chance level corresponds to the success rate expected by random target selection. Shared-control gain is the difference in success rate between shared-control and BCI-only conditions, expressed in percentage points (pp). The last column reports the p-value from a paired, two-sided Wilcoxon signed-rank test across sessions, testing whether success rates differed between shared-control and BCI-only conditions.

| Monkey | Task | Sessions | BCI-only (%) | Shared-control (%) | Chance (%) | Gain (pp) | *p* |
| --- | --- | --- | --- | --- | --- | --- | --- |
| Monkey 1 | Fixed Obstacle | 12 | 49 ± 10 | 81 ± 8 | 14 | 31.6 | 4.88 × 10<sup>-4</sup> |
| Monkey 1 | Appearing Obstacle | 10 | 39 ± 15 | 67 ± 16 | 18 | 27.4 | 1.95 × 10<sup>-3</sup> |
| Monkey 1 | Appearing Obstacle (6 months later) | 9 | 57 ± 12 | 80 ± 10 | 16 | 23.0 | 3.91 × 10<sup>-3</sup> |
| Monkey 1 | Respawn | 9 | 79 ± 8 | 67 ± 10 | 24 | -12.6 | 1.17 × 10<sup>-2</sup> |
| Monkey 2 | Fixed Obstacle | 11 | 34 ± 8 | 61 ± 9 | 18 | 27.1 | 9.77 × 10<sup>-4</sup> |
| Monkey 2 | Appearing Obstacle | 9 | 22 ± 9 | 56 ± 9 | 21 | 33.7 | 3.91 × 10<sup>-3</sup> |

**Extended Data Table 3.** Comparison of fixed-blending and prior-only control in the brain-computer interface (BCI)-like Fixed Obstacle environment. Performance across 100 matched simulated trials for user-only control, fixed linear blending with different prior weights \(W\), prior-only control, and full shared control. The BCI-like environment reproduced the five-target geometry of the monkey Fixed Obstacle task with a single static obstacle positioned between the start position and designated target. Success, collision, and timeout percentages sum to 100% within each controller condition. The highest success rate among the tested fixed-blending weights occurred at W=0.75

| Controller | Parameter | Success (%) | Collision (%) | Timeout (%) |
| --- | --- | --- | --- | --- |
| User-only | – | **0** | 100 | 0 |
| Fixed blend | W = 0.25 | 0 | 100 | 0 |
|  | W = 0.50 | 6 | 94 | 0 |
|  | W = 0.75 | **88** | 12 | 0 |
|  | W = 0.80 | 82 | 0 | 18 |
|  | W = 0.85 | 81 | 0 | 19 |
|  | W = 0.90 | 79 | 0 | 21 |
|  | W = 0.95 | 46 | 0 | 54 |
| Phase-specific | R = 1.5 m | 12 | 88 | 0 |
|  | R = 2.0 m | **100** | 0 | 0 |
|  | R = 2.5 m | 100 | 0 | 0 |
|  | R = 3.0 m | 100 | 0 | 0 |
| Prior-only | – | **28** | 0 | 72 |
| Full shared control | – | **97** | 0 | 3 |

**Extended Data Table 4.** Comparison of fixed-blending and prior-only control in the randomized cluttered simulation environment. Performance across 100 matched simulated trials for user-only control, fixed linear blending with different prior weights W, prior-only control, and full shared control. The randomized environment contained 1–6 candidate targets and 1–6 static obstacles with scene configurations varying across trials. Success, collision, and timeout percentages sum to 100% within each controller condition. The highest success rate among the tested fixed-blending weights occurred at W=0.85.

| Controller | Parameter | Success (%) | Collision (%) | Timeout (%) |
| --- | --- | --- | --- | --- |
| User-only | – | **26** | 74 | 0 |
| Fixed blend | W = 0.25 | 31 | 69 | 0 |
|  | W = 0.50 | 37 | 63 | 0 |
|  | W = 0.75 | 56 | 44 | 0 |
|  | W = 0.80 | 63 | 36 | 1 |
|  | W = 0.85 | **73** | 19 | 8 |
|  | W = 0.90 | 61 | 6 | 33 |
|  | W = 0.95 | 34 | 1 | 65 |
| Phase-specific | R = 1.5 m | 42 | 53 | 5 |
|  | R = 2.0 m | **73** | 3 | 24 |
|  | R = 2.5 m | 68 | 2 | 30 |
|  | R = 3.0 m | 63 | 2 | 35 |
| Prior-only | – | **17** | 0 | 83 |
| Full shared control | – | **75** | 4 | 21 |

